# A generalizable normalization framework to decouple protocol and instrument effects: Application to high-sensitivity proteomics multicentric study (PME13)

**DOI:** 10.64898/2026.08.16.744113

**Authors:** Gianluca Arauz-Garofalo, Sergio Ciordia, Anne Gonzalez de Peredo, Karima Chaoui, Jeewan Babu Rijal, Valeriane Gaxotte, Ignasi Folch-i-Casanovas, Mikel Azkargorta, Ruben Almey, Kerman Aloria, Büşra Aytül Kirim, Rodrigo Barderas, Sophie Braga-Lagache, Enrique Calvo, Eduardo Chicano-Galvez, Felipe Clemente, Gabriela N. Chirițoiu, Cristina Chiva, Mathilde Decourcelle, Maarten Dhaenens, Ramón Díaz-Peña, Thibaut Douché, Alba Durán-Cortines, Mª Carmen Durán-Ruiz, Khadija El Koulali, Almudena Escobar-Niño, Francisco Javier Fernandez Acero, Jokin Fernández-Irigoyen, Carlos García-García, Concha Gil, Sandra Goetze, Eloisa Gonzalez Vidal, MD Gutierrez, María Luisa Hernáez, Cristina María López, Consuelo Marín-Vicente, María Luisa Mateos-Martín, Mariette Matondo, Ana Montero-Calle, Esperanza Morato-López, Cristian V.A. Munteanu, Antonia Odena-Caballol, Ignacio Ortea, Nazlı Ezgi Özkan, Nurhan Özlü, Ángela Peralbo-Molina, Davide Perico, Vivian De los Ríos, Eduard Sabidó, María Luz Valero, Bart Van Puyvelde, Jesús Vázquez, Raffaello Viganò, Mar Vilanova, Marta Vilaseca, Simon D. Widmer, Ines Zapico, Camille Stephan-Otto Attolini, Ferdinando Cerciello, Christine Carapito, Odile Burlet-Schiltz, Felix Elortza, Marina Gay, Fernando J. Corrales

## Abstract

Multicenter studies are essential for benchmarking analytical workflows, yet their interpretation is often confounded by the combined effects of experimental protocols and instrumentation. To address this challenge, we introduce a simple normalization-based analytical framework, the recovery metric (*ρ*), designed to decouple protocol driven effects from instrument dependent variability.

We applied this framework to the 13^th^ Proteomics Multicentric Experiment (PME13), a large multicentric proteomics dataset generated across 27 laboratories using high sensitivity workflows and varying sample preparation protocols. By leveraging a common digested reference sample, *ρ* enables direct cross-comparison of all datasets on a unified scale, effectively minimizing instrument-related biases.

Using this approach, we demonstrate that apparent instrument dependent trends are largely removed when evaluated through *ρ*, revealing consistent protocol driven effects across laboratories. Statistical modeling identified key variables influencing *ρ*, including sample input amount, reduction and alkylation, and the use of n-dodecyl-β-D-maltoside (DDM). While DDM was associated with improved *ρ*, reduction and alkylation and additional handling steps led to reduced performance, particularly at low input levels.

We further highlight practical considerations for the application of ratio based normalization, including the occurrence of values exceeding theoretical bounds, which reflect deviations from underlying assumptions and require appropriate filtering.

Overall, this work establishes a generalizable analytical strategy for disentangling confounding factors in multicentric datasets and provides practical guidelines for optimizing high sensitivity proteomics (HSP) workflows. The proposed framework may be broadly applicable to other analytical fields where cross laboratory comparability is required.

## Introduction

Continuous advances in proteomics over the last two decades have been driven by improvements in mass spectrometry instrumentation, sample preparation workflows, and bioinformatics tools. Together, these developments now enable near-complete profiling of complex proteomes in a single liquid chromatography coupled to mass spectrometry (LC-MS/MS) run [1]. In particular, optimized sample preparation protocols are critical to ensure high peptide yield, reproducibility, and accurate quantification, especially in high-sensitivity proteomics (HSP) applications.

The rapid development of single-cell proteomics (SCP) has further emphasized the need for highly efficient and reproducible workflows. Understanding the complexity of a tissue depends on defining the contributions of individual cells rather than relying on averaged analyses of the whole entity, as demonstrated by single-cell RNA sequencing studies [2]. However, transcriptomic profiling alone is insufficient; complementing it with proteomic analysis is essential to accurately characterize cellular phenotypes, as proteins serve as the primary functional effectors within the cell. Recent advances in instrumentation and workflow miniaturization have enabled the identification of thousands of proteins from extremely low input samples [3,4], supported by improvements in sample processing [5,6], chromatography [7], mass spectrometry acquisition and quantification [8–10], and data analysis strategies [11–13].

Despite these advances, establishing robust and transferable workflows for low-input proteomics remains challenging. Multicenter studies are essential to benchmark protocols, define best practices, and ensure reproducibility across laboratories [14]. Large collaborative initiatives, including Human Proteome Organization Proteomics Standards Initiative (HUPO-PSI) [15], Association of Biomolecular Resource Facilities (ABRF) [16], as well as the Clinical Proteomic Tumor Analysis Consortium (CPTAC) [17], have made substantial contributions toward standardization, data sharing, and quality control. In the same direction, the Spanish consortium ProteoRed consolidated an interlaboratory-based standardization initiative [18–22], which was later expanded to a European dimension as the EuPA Standardization Initiative, aiming to promote a common space integrating proteomic strengths across Europe, in order to increase the penetrance of proteomics in other scientific disciplines and to facilitate the implementation of state of the art methods and technology in emergent platforms under standardized conditions [23]. All these efforts underscore the importance of setting protocol benchmarking and quality control evaluation principles, as well as valuable reference samples to facilitate the adoption of emerging technologies on less experienced platforms, ensuring competitive performance and promoting the expansion of proteomics across various scientific domains.

However, the interpretation of multicenter proteomics studies focused on sample preparation protocols remains fundamentally limited by the confounding effects between protocol-driven and instrument-driven variability. Differences in instrument performance can mask or exaggerate the effects of sample preparation, making it difficult to derive robust and transferable conclusions [24].

To address this challenge, we propose a simple and generalizable normalization framework designed to decouple sample preparation effects from instrument-dependent variability. By leveraging a common reference sample, this approach enables meaningful cross-laboratory comparisons and provides a robust basis for evaluating protocol performance in multicentric studies.

To demonstrate the utility of this framework, we conducted the 13^th^ Proteomics Multicentric Experiment (PME13) within the EuPA Standardization Initiative, focusing on high-sensitivity proteomics (HSP). This interlaboratory study provides a unique opportunity to evaluate the performance and reproducibility of HSP workflows across laboratories, platforms, and sample preparation strategies, while establishing practical guidelines for protocol optimization.

## Materials and methods

### Study design

We selected a standardized *E. coli* extract as the benchmarking sample for PME13 (BioRad, Ref. 163-2110). Two different samples were prepared from the same *E. coli* protein extract: one sample with an undigested protein extract (from now on, high sensitivity sample or “HS”) and one sample with an already digested protein extract (from now on, reference sample or “Ref”). Establishing these two sample typologies will be key to decouple protocol and instrument effects if a derived quantification magnitude is properly defined. Our proposal for such quantification magnitude will be the recovery metric *ρ* (see “Recovery definition and inferential analysis” subsection for further details). By leveraging this *ρ* we will enable the comparison of distinct sample preparation protocols across independent laboratories.

### Sample preparation and LC-MS analysis

For the HS (undigested) sample, each vial containing 2.7 mg of *E. coli* protein extract was resuspended in 100 mM triethylammonium bicarbonate (TEAB) to achieve a concentration of 500 ng/µL (HS initial stock). To prepare the Ref (digested) sample, 10 tubes containing 80 µg of *E. coli* protein extract in sodium dodecyl sulfate (SDS) buffer were prepared. These were reduced and alkylated separately with 5 mM tris(2-carboxyethyl)phosphine (TCEP) and 10 mM chloroacetamide (CAA), respectively, and digested on S-Trap columns with trypsin (1:15 enzyme-to-protein ratio). The samples were dried and directly resuspended in buffer A containing 2% ACN and 0.1% FA. Peptide quantification was performed using fluorescence with Qubit™. Once similar peptide recovery was confirmed, they were combined into a pooled stock at a concentration of 500 ng/µL (Ref initial stock). From these two initial stocks (HS and Ref), and using the Opentrons OT-2 robot, three serial dilutions (1:10, v/v) were prepared with 25 mM TEAB to obtain final stocks of 5, 0.5, and 0.05 ng/µL. 20 µL of each stock were dispensed into 8-well strips and dried in a Speed-Vac to prepare study samples containing total amounts (or tiers) of 100, 10, and 1 ng.

Each PME13 participant received 4 lyophilized pseudo-biological replicates of *E. coli* per tier (half a strip), both for HS and Ref samples (24 samples in total) for the mandatory data dependent acquisition (DDA) characterization. Additional sample sets were provided to those laboratories willing to also generate an optional data independent acquisition (DIA) characterization. Basic analysis guidelines (**Supplementary Document 1**), like the standardized spectrum file naming and injection order to be followed (by increasing tier order, 1 ng - 10 ng - 100 ng, always with the HS replicates quartet preceding the Ref replicates quartet within each given tier) were also shared with all laboratories.

Each participating laboratory employed its own preferred LC-MS/MS configuration. Importantly, within each dataset, the same LC-MS/MS setup and acquisition parameters were consistently applied to both the HS and Ref samples, to meet the requirements of the downstream recovery framework used to unlock a reliable cross-laboratory comparison. A comprehensive overview of all LC-MS/MS systems is provided in the supplementary metadata table (**Table S1**). The mass spectrometry proteomics data have been deposited to the ProteomeXchange Consortium via the Proteomics Identifications Database (PRIDE) [25] partner repository with the dataset identifier PXD082645. After sample analysis, participants returned the raw data or spectrum files and detailed metadata information regarding the most relevant sample preparation conditions (reduction, alkylation, tube transference, digestion enzyme, buffer, etc) and the LC-MS/MS instrumentation used to generate the MS data (HPLC, gradient, MS instrument, MS2 detector, etc).

### Data analysis

The workflow implemented in this study is illustrated in **Figure 1**.

**Figure 1.**
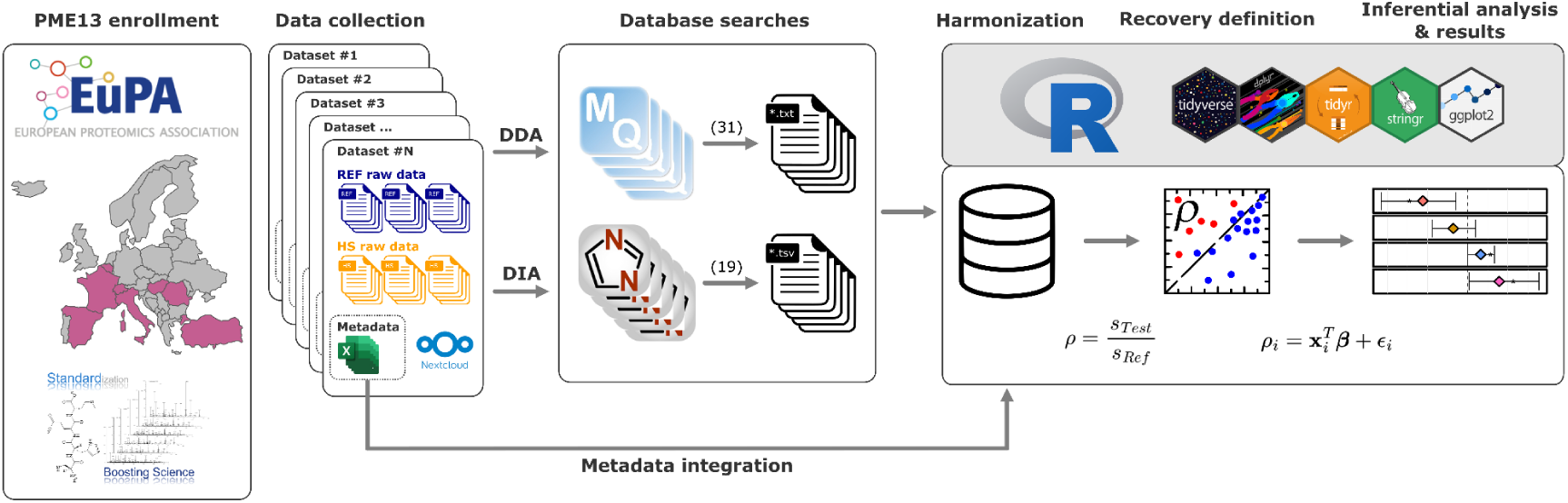
Diagram of the workflow and the analysis procedure followed in this study. The process includes enrollment to the PME13; sample delivery; spectrum file collection, HS and Ref samples generated via DDA and (optionally) DIA; centralized database searching; data harmonization and integration; data visualization and analysis; and finally the generation of results. (PME13: 13^th^ Proteomics Multicentric Experiment; Ref: Reference; HS: High sensitivity; DDA: Data dependent acquisition; DIA: Data independent acquisition).

#### Data collection

We collected the raw LC-MS/MS files along with their associated metadata spreadsheets via NextCloud. The metadata included the following information divided in four main categories:

**General information:** Laboratory identifier, Method, Starting amount (ng), Injected amount (ng), Spectrum file name.

##### Sample preparation

● Tube transfer: TRUE/FALSE depending on whether the sample was transferred to a different tube during the experimental procedure.
● Reduction: TRUE/FALSE depending on whether disulfide bonds were reduced.
● Alkylation: TRUE/FALSE depending on whether disulfide bonds were alkylated after reduction.
● Alkylation Reagent: Alkylating agent used.
● Enzyme: Enzyme used in the digestion process.
● Enzyme amount (ng): Amount of enzyme used.
● Digestion Buffer: Solution in which digestion was carried out.
● Volume of digestion buffer (µL): Digestion volume.

##### LC-MS/MS system

● High-performance liquid chromatography (HPLC): Liquid chromatography instrument used.
● Gradient (min): Gradient time used in chromatographic separation.
● Length (cm): Chromatographic column length.
● MS instrument: Model of the mass spectrometer.
● MS2 detector: Type of MS2 detector used.

**Notes:** Free text field to add any comments considered relevant.

#### Database searches

We performed centralized database searches using two different software tools: MaxQuant (v2.4.2.0) [26] for DDA datasets and DIA-NN (v1.9.2) [26,27] for DIA datasets. In both cases, the same databases were used for the search: SwissProt for *E. coli* and *H. sapiens* (May 2023), plus a universal contaminants database [28]. All search parameter files are available at PRIDE (PXD082645) [25].

We searched each dataset independently, with the premise of sticking as closely as possible to default parameters, disabling match between runs, adding methionine oxidation as a dynamic modification and carbamidomethylation on cysteine only when reduction and alkylation had been performed in the experimental procedure. For DIA-NN, we enabled the “Heuristic protein inference” option and selected “Protein name (from FASTA),” as well as “QuantUMS (high precision)” for the quantification strategy. For DIA dataset ID85, DIA-NN parameters were adapted to accommodate the MS2 acquisition performed on a linear ion trap [29].

#### Harmonization of peptide and protein identifications and metadata integration

All data analyses were performed in R using a range of packages, including ggplot2, tidyverse-related tools, and other statistical and visualization libraries (e.g., lme4, pheatmap, MASS).

Output files from MaxQuant and DIA-NN differ substantially in structure and content. The *evidence.txt* file from MaxQuant and the *report.parquet* file from DIA-NN were selected, as they provide the most comprehensive information. We filtered FDR 1 % at peptide and protein levels. Next, we also dropped those proteins identified with only one peptide within each dataset.

Given the lack of standardization across proteomics search engines, a subset of relevant variables was selected and unified across datasets (e.g., sequence, protein group, intensity, annotations). Column names were standardized regardless of the originating software, and additional variables were introduced, including the source software, protein taxonomy, and a contaminant flag. Metadata and identification results were then integrated into a single table using the file identifier (**Table S1**).

Depending on the focus, data were summarized at multiple levels (method, dataset, sample, input amount or lab). Quantitative metrics included counts of PSMs/precursors, peptides, protein groups (PGs) and normalized intensity values (normalization by total intensity per sample and scaling to the maximum total intensity observed across all replicates).

Coefficients of variation (CVs) were calculated using the standard deviation and mean normalized intensities of each PG across replicates within the same dataset, method, and sample input.

#### Recovery definition and inferential analysis

To account for the effects of instrument and acquisition method, we propose the recovery metric (*ρ*). This *ρ* was defined as the ratio between the signals of a testing sample (*s*_Test_) and a reference sample (*s*_Ref_):

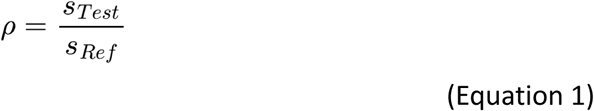

Note that *ρ* definition is general once Test and Ref sample types are established. In the proteomics arena, the “signals” appearing in **Equation 1** could be any quantifiable magnitude characterizing the information yield obtained from Test and Ref samples, for example: the intensity or count of identified peptide spectrum matches (PSMs), peptides, PGs or precursors.

In the particular case of our HSP multicentric study, for the DDA dataset we pick up the PSM median count in HS (Test in **Equation 1**) and Ref samples as **Equation 1** signals, respectively. Analogously for the DIA dataset, we pick up the Precursor median count in HS and Ref samples as **Equation 1** signals. Therefore, we worked with the following two rho definitions:

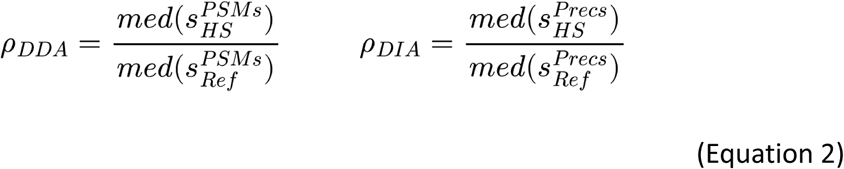

On the one hand, in the DDA dataset, working with PSM counts instead of precursor abundances partially mitigates the negative effect of losing information for identified-but-not-quantified peptides. For symmetry, we decided to work with Precursor counts in the DIA dataset (from now onwards, we will use “features” to name PSMs or precursors in DDA or DIA datasets, respectively). On the other hand, working with features offered greater resolution and granularity, allowing for more refined data evaluation compared to a peptide or PG level recovery metric. Finally, using the replicate medians for feature counts aggregation offered a more stable measure when compared to other options, such as maximums or means. Keep in mind that, in other omics scenarios like metabolomics or transcriptomics, the signals to be plugged in **Equation 1** as s_Test_ and s_Ref_ could be chosen at will depending on the particularities of each dataset.

A linear model was devised to assess the impact of sample preparation variables on *ρ*. We applied a Box-Cox transformation (*λ* = 0.58) to meet the assumptions of the linear model [30]:

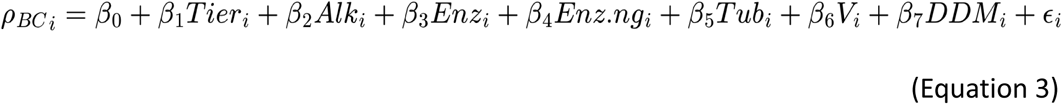

where *ρ*_BC_ (the Box Cox transformed *ρ*) is the response variable, *ε* is the error term, and the *β_j_* (*j* = 1, …, 7) are the regression coefficients of each explanatory variable used, namely: "Tier" (log_10_ of the starting sample amount in ng), “Alk” (boolean specifying if reduction and alkylation was applied), “Enz” (categorical specifying the digesting enzyme, Trypsin or Trypsin + LysC), “Enz.ng” (digesting enzyme amount in ng), “Tub” (boolean specifying if sample tube transfer took place), “V” (digestion buffer volume in μL) and “DDM” (boolean specifying if DDM was applied).

## Results and discussion

### PME13 participation summary and overall identifications overview

Samples were distributed to 35 laboratories across Europe. From those, 27 laboratories, from 8 different countries, returned data, Spain being the country with the highest number of participants (**Figure S1**). Almost half of the laboratories (17 out of 35) had not participated in previous PMEs. A total of 50 datasets were collected (62% DDA).

We processed a total of 1200 raw files from the 50 datasets, 31 DDA and 19 DIA (∼1.6 TBs). Overall, across all datasets and both acquisition methods, we identified a total of 1,630,477 features (PSMs for DDA and precursors for DIA) and 40,495 peptides corresponding to 2,937 *E. coli* PGs. As expected, the number of *E. coli* identifications drops at all levels (PGs, peptides, and features) as the amount of input material decreases. For instance, on average the number of *E. coli* PGs doubles from 1 to 10 ng and increases by 38% from 10 to 100 ng (607, 1,354, and 1,867 *E. coli* PGs in 1, 10, 100 ng tiers, respectively). DIA identifications were higher than DDA, despite having 63% fewer datasets (31 vs. 19). We identified 1,292 *E. coli* PGs common to both methods, with 557 PGs unique to DDA and 1,054 PGs unique to DIA. Specifically, we observed 26% more PGs and 27% more peptides identified from *E. coli* in DIA datasets. Contrary to our expectations, in all but one DDA dataset, we detected at least one *E. coli* PSM in some replicates of the 1 ng tier, while only 48% of the DIA datasets detected at least one precursor at 1 ng tier. Identifying even a single protein from as little as 1 ng of starting material is a considerable challenge, especially for older mass spectrometers, such as the LTQ Orbitrap Velos Pro (Thermo). On the other hand, we found that 607 *E. coli* PGs were identified in 50% of DDA datasets, and 1,034 PGs in the case of DIA (**Figure 2**).

**Figure 2.**
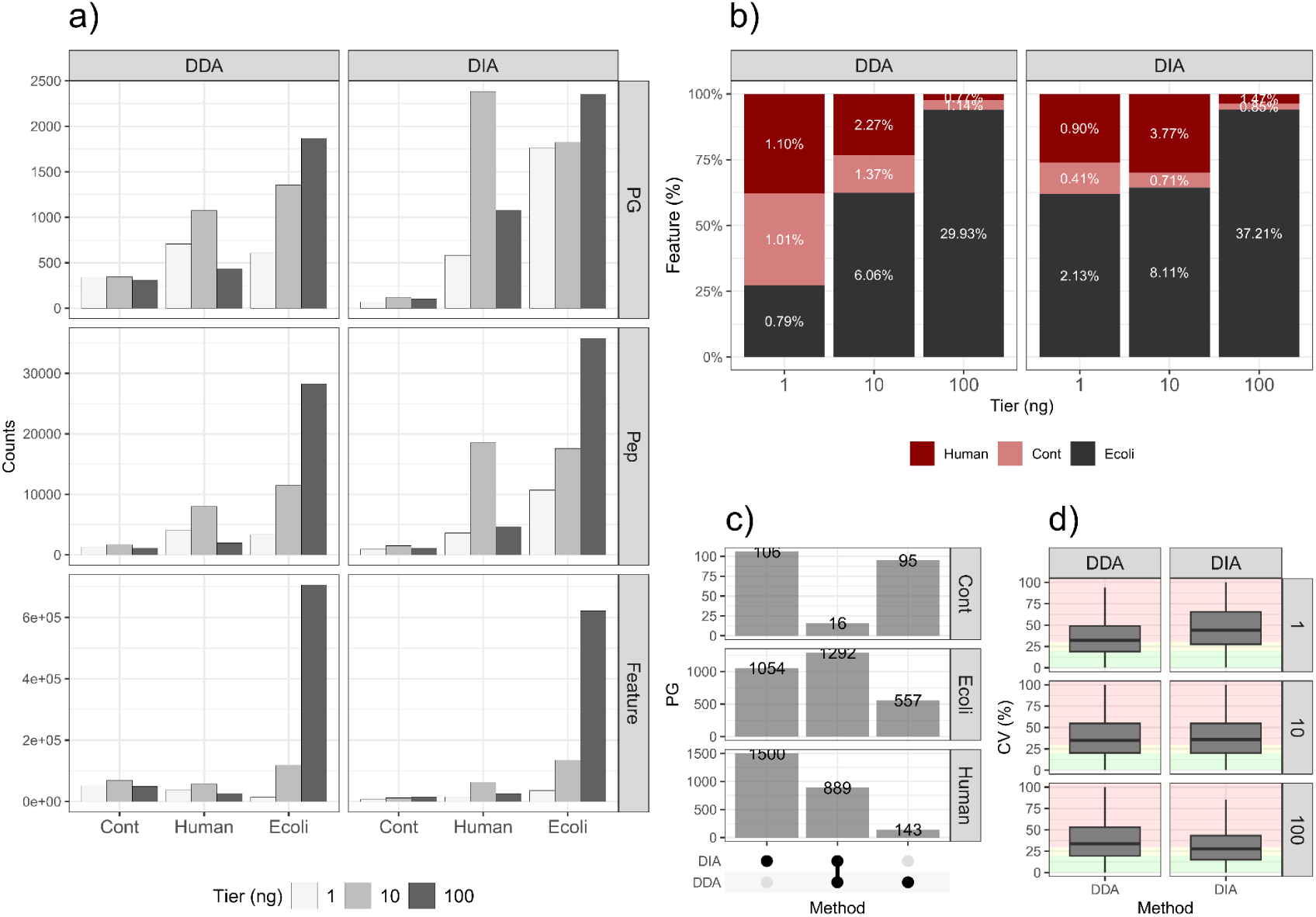
Global identifications and CVs. **a)** Bar chart showing the number of identifications (y-axis) per database (x-axis) for PGs, Peps, and PSMs or precursors (Features) (grid rows), and for each acquisition method (grid columns). The gray shade represents the initial sample amount. **b)** Stacked bar chart representing the percentage of features (y-axis) as a function of the initial sample amount in ng (x-axis) for each acquisition method (grid columns). Global percentages within each acquisition method are shown as labels and color differentiates the database. **c)** Upset plot with the number of PGs (y-axis) for all possible set intersections between DDA and DIA (x-axis), distinguishing the database in which the identification was made (grid columns). Absolute PG counts for each intersection are also indicated (labels). **d)** Boxplots showing the CVs (outliers not shown), expressed as a percentage, for each PG (y-axis) differentiating the acquisition method (x-axis and grid columns) and the tier (grid rows). Background colors represent CV ranges: green: CV < 20%, yellow: 20% < CV < 30%, red: CV > 30%. (CV: Coefficient of variation; PG: Protein group; Pep: Peptide; PSM: Peptide spectrum match; DDA: Data dependent acquisition; DIA: Data independent acquisition; Cont: Contaminant).

Apart from the *E. coli* PGs we detected a total of 2,527 peptides and 498 PGs from the contaminant database. We also identified 2,749 PGs, derived from 21,841 peptides and 223,164 features from the *H. Sapiens* protein database. The total number of human PGs is nearly equal to that of *E. coli*. When breaking down this information by tier and acquisition method, we observe that the number of human or contaminant PGs are generally high but still lower than those from *E. coli*. However, in some cases, peptide and feature identifications from human proteins exceed those from *E. coli*, suggesting that some contaminant proteins may be highly abundant. For peptides and features, there is a clear trend: the proportion of human and contaminant identifications increases relative to *E. coli* as the sample input amount decreases. This effect is clearer in DDA than in DIA (**Figures 2a and 2b**). We also observed that DIA yielded many more unique human identifications than DDA (**Figure 2c**).

We then examined the number of identifications in each replicate and dataset of the HS sample, differentiating identifications coming from *E. coli* or contaminants (defined here as any identification from *H. sapiens* or contaminant databases). We observed that the number of contaminants was dataset and laboratory specific. In other words, most contaminants came from a few datasets (ID28, ID72, ID62, ID95), where *E. coli* samples had likely become contaminated either during sample preparation or in the chromatographic column itself, due to analytical column carry-over from previous runs (**Figure S2**).

All these observations highlight that using *E. coli*, rather than human material, as the standard sample for HSP was indeed a good choice. It allows us to monitor contamination levels and emphasizes the need to follow contamination free protocols when working with very low protein quantities, especially when working with human samples.

Next, we calculated the coefficient of variation (CV) for each PG across the four replicates to evaluate the reproducibility. In general, CVs were high, in most cases with medians above 30%. As expected, CVs increased as the amount of starting material decreased. At 100 ng and 10 ng, DIA had slightly lower CVs than DDA. Interestingly, at 1 ng, DDA showed lower CVs than DIA (**Figure 3d**). At first glance, this insight appears to be heavily influenced by dataset ID36, which at 1 ng had the highest CVs and also the highest identification numbers of the whole PME13 (**Figure S3**). However, this trend persisted even after excluding ID36, with DIA median CVs still exceeding those of DDA. In our study, the widely accepted superiority of DIA over DDA [31], seems to vanish at low input levels for the percentage of laboratories identifying *E.coli* proteins and especially the CV quality. One possible explanation is that DIA may offer less benefit against DDA when decreasing sample complexity.

**Figure 3.**
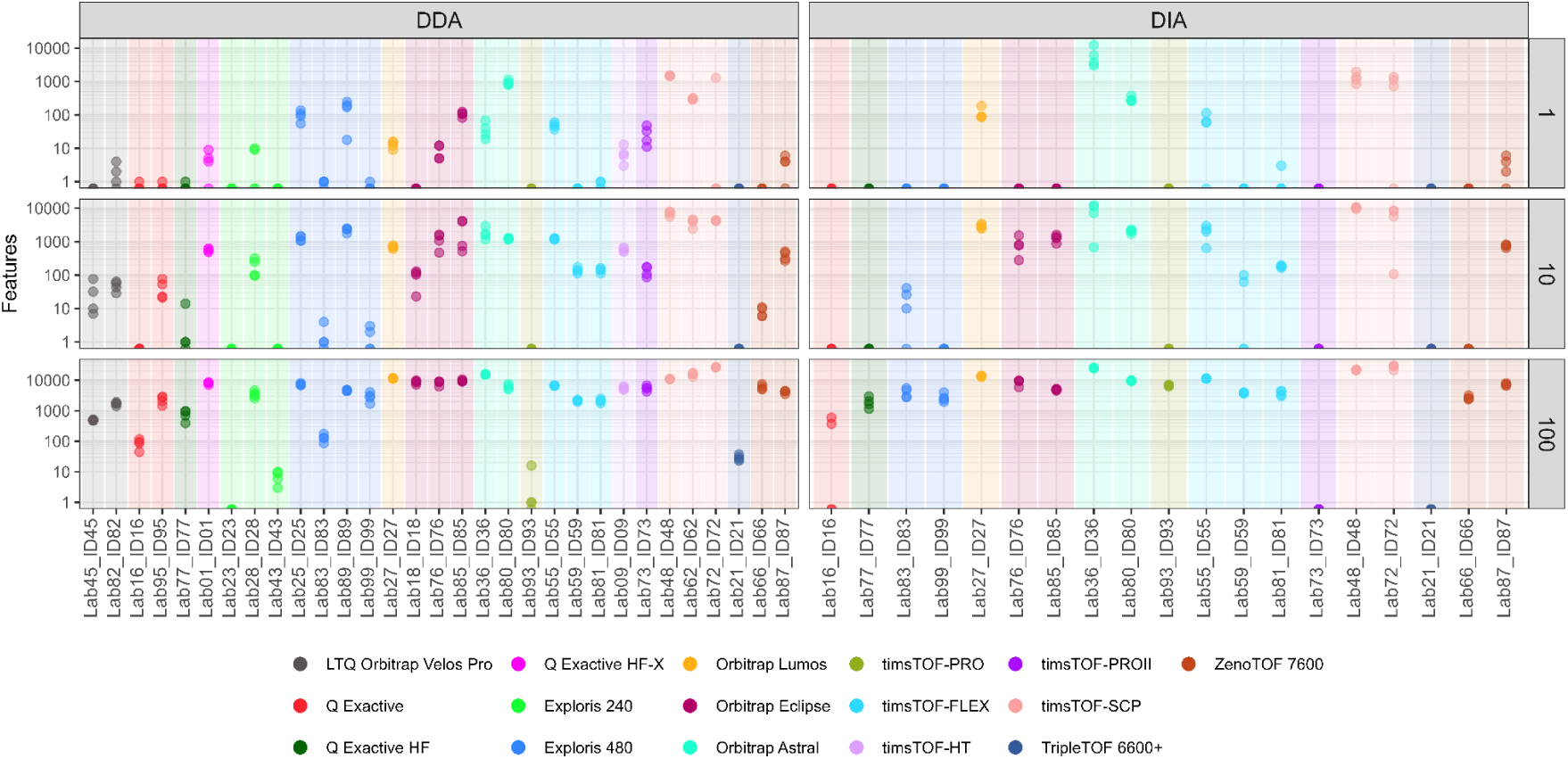
Identification numbers by dataset highlighting mass spectrometers. Stripcharts showing the number of features (y-axis, in log_10_ scale) across laboratories and datasets (x-axis). The colors differentiate the mass spectrometers used. Grid rows indicate the initial sample amount, and grid columns the acquisition method. The features shown in the left column are PSMs and those in the right column are precursors, corresponding to DDA and DIA, respectively. (PSM: Peptide spectrum match; DDA: Data dependent acquisition; DIA: Data independent acquisition).

### Instrument impact and instrument-wise qualitative analysis

One of the main challenges we encountered is the strong dependence between each laboratory performance and the mass spectrometer used. Newer and more expensive instruments should yield better results. To confirm this, we visualized the data according to instrumentation (**Figure 3**). We observed substantial differences in the number of features identified based on the instrument used, but we also noticed that instrumentation alone does not explain all performance variability. In fact, dissimilar results were obtained even between datasets using the same instrument. For this reason, we decided to carry out a streamlined and separate analysis for those instruments with multiple datasets available.

We performed a qualitative, manually curated analysis, with the intention of identifying general trends consistent across instruments. We evaluated the outcome metrics for each dataset available within a given instrument, looking for patterns in concordance with their corresponding metadata. We used all available metrics and samples (HS and Ref), i.e., we visualized features, peptides, and PGs for both the HS and Ref samples at 1, 10, and 100 ng, across the four available replicates.

Following this qualitative evaluation, we identified general trends in experimental protocols that could explain the outcome disparity (**Figure S4**). In particular, we observed that the use of commercial kits and/or extra clean-up steps seem to hinder protein detection, likely due to added sample loss during the process. In contrast, the use of Field Asymmetric Ion Mobility Spectrometry (FAIMS) enhances protein identification, particularly at low input concentrations.

### Recovery metric and inferential analysis of experimental variables

A central challenge in multicentric studies is separating protocol performance from instrument-driven variability. To address this, we defined the *ρ* framework and used it to compare datasets on a common scale (**Equation 2**). Within this framework, using PSM counts rather than precursor intensities in *ρ* definition, avoided the exclusion of a noticeable number of identified-but-not-quantified precursors from *ρ* computation, namely 8%, 6%, and 4% at input levels of 1, 10, and 100 ng, respectively. We found that at 100 ng, most *ρ* values approached 1, denoting optimal performance for a big majority of the datasets available. Nevertheless, at 1 ng, *ρ* values dropped significantly, even reaching zero for some datasets (**Figure 4a**). Instrument-dependent trends previously observed were no longer apparent when using *ρ*, confirming its ability to decouple sample preparation performance from instrument effects (**Figure 4b**). For example, two datasets acquired on a different instrument generation, ID62 (timsTOF-SCP) and ID55 (timsTOF-FLEX), showed markedly different identification counts at tier 1 ng for DDA (301 vs. 48 PSMs, respectively), yet yielded very similar *ρ* values (0.89 and 0.85). This suggests that both sample preparation protocols of ID62 and ID55 performed effectively, notwithstanding the disparate capabilities of the instruments used to generate the datasets.

**Figure 4.**
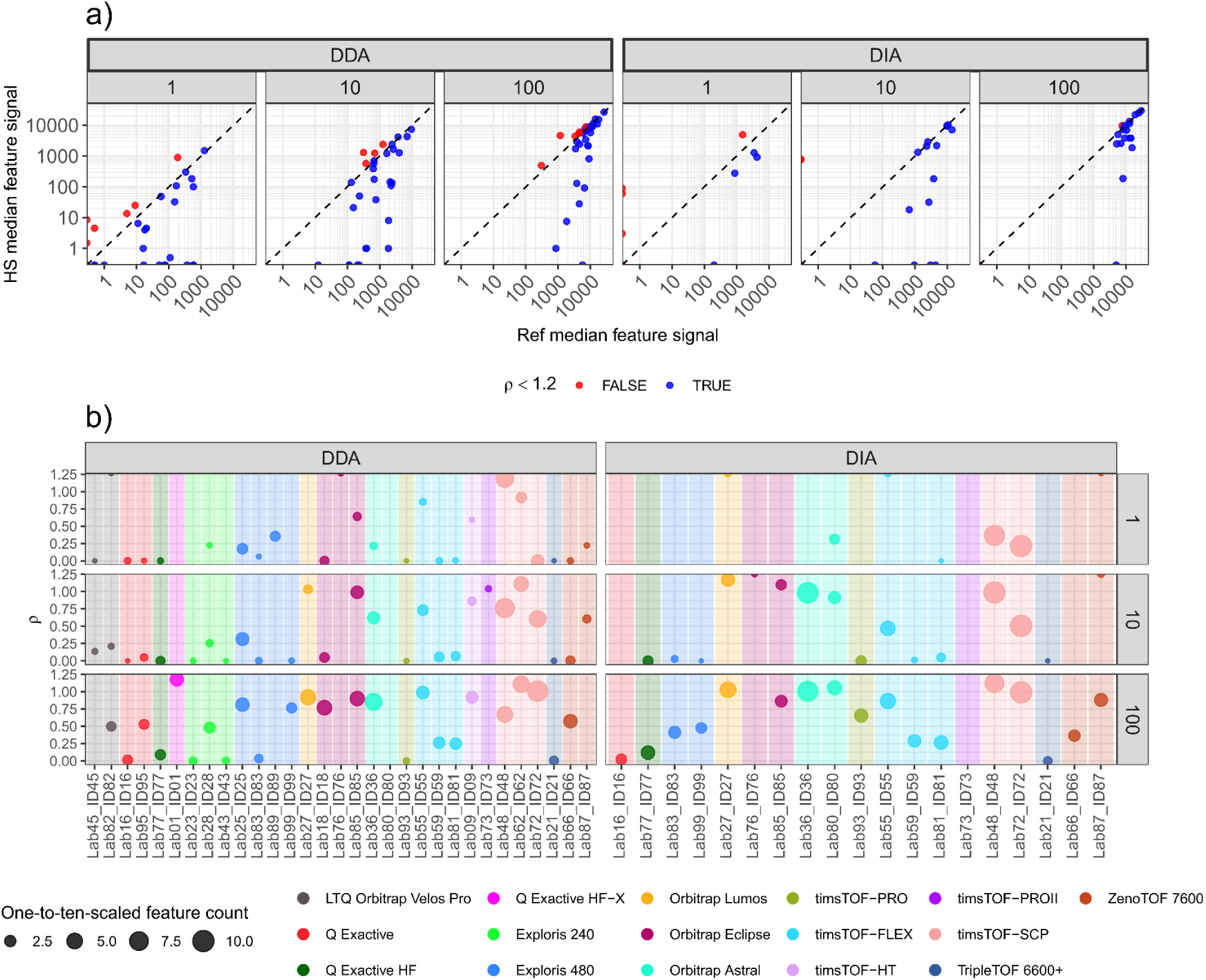
Recovery metric definition and overview. **a)** Scatter plot with the HS sample median feature signal (y-axis, in log_10_ scale) versus the Ref sample median feature signal (x-axis, in log_10_ scale), for each acquisition method and each initial sample amount (nested grid columns). The oblique dashed line represents the identity, with a slope 1 and intercept 0. Colors indicate whether *ρ* is considered valid according to our threshold (*ρ* < 1.2). **b)** Dot plot showing *ρ* (y-axis) for each laboratory and dataset (x-axis), differentiating the mass spectrometer (color). The initial sample amount is also represented (grid rows) along with the acquisition method (grid columns). The size of the dots corresponds to the number of features used to compute *ρ* (one-to-ten scaled within each input amount), which are PSMs for DDA and precursors for DIA. (Ref: Reference; HS: High sensitivity; *ρ*: Recovery metric; PSM: Peptide spectrum match; DDA: Data dependent acquisition; DIA: Data independent acquisition).

Values of *ρ* greater than 1 indicate violations of the underlying assumption that the reference sample represents an upper bound of recovery. These cases likely arise from technical inconsistencies (e.g., incomplete resuspension of the reference sample) and were therefore excluded using a predefined threshold. We accepted a 20% tolerance and discarded all samples with *ρ* > 1.2 from the downstream inferential analysis to prevent misinterpretation (retaining 73% of the original dataset) (**Figure 4a**).

To identify experimental variables influencing *ρ*, we fitted a linear model (**Equation 3**). Rather than aiming to accurately predict *ρ*, this model was designed to capture global trends and assess the relative impact of sample preparation variables. The model should therefore be interpreted as an exploratory tool to identify robust trends rather than causal effects. The analysis identified three significant factors: reduction and alkylation, DDM, and input amount (tier) (**Table 1 and Figure 5**). No significant associations were observed for enzyme choice or tube transfer. Although tube transfer was expected to negatively affect *ρ* due to the additional sample manipulation and the associated risk of sample loss, the estimated coefficient showed a slight and non-significant increase in *ρ* when tube transfer was performed. Similarly, enzyme choice showed a non-significant tendency toward lower *ρ* with Trypsin+LysC compared with Trypsin alone. Enzyme amount and digestion buffer volume were also non-significant, with coefficient estimates close to zero, suggesting no detectable association with *ρ* under the conditions evaluated. To highlight the significant effects, we examined *ρ* values across Tier (**Figure 5**). Reduction and alkylation was consistently associated with *ρ* reduction across all input levels, whereas the use of DDM led to a consistent improvement in *ρ*. Additionally, *ρ* is reduced with decreasing input amount, confirming the expected relationship between sample quantity and recovery.

**Figure 5.**
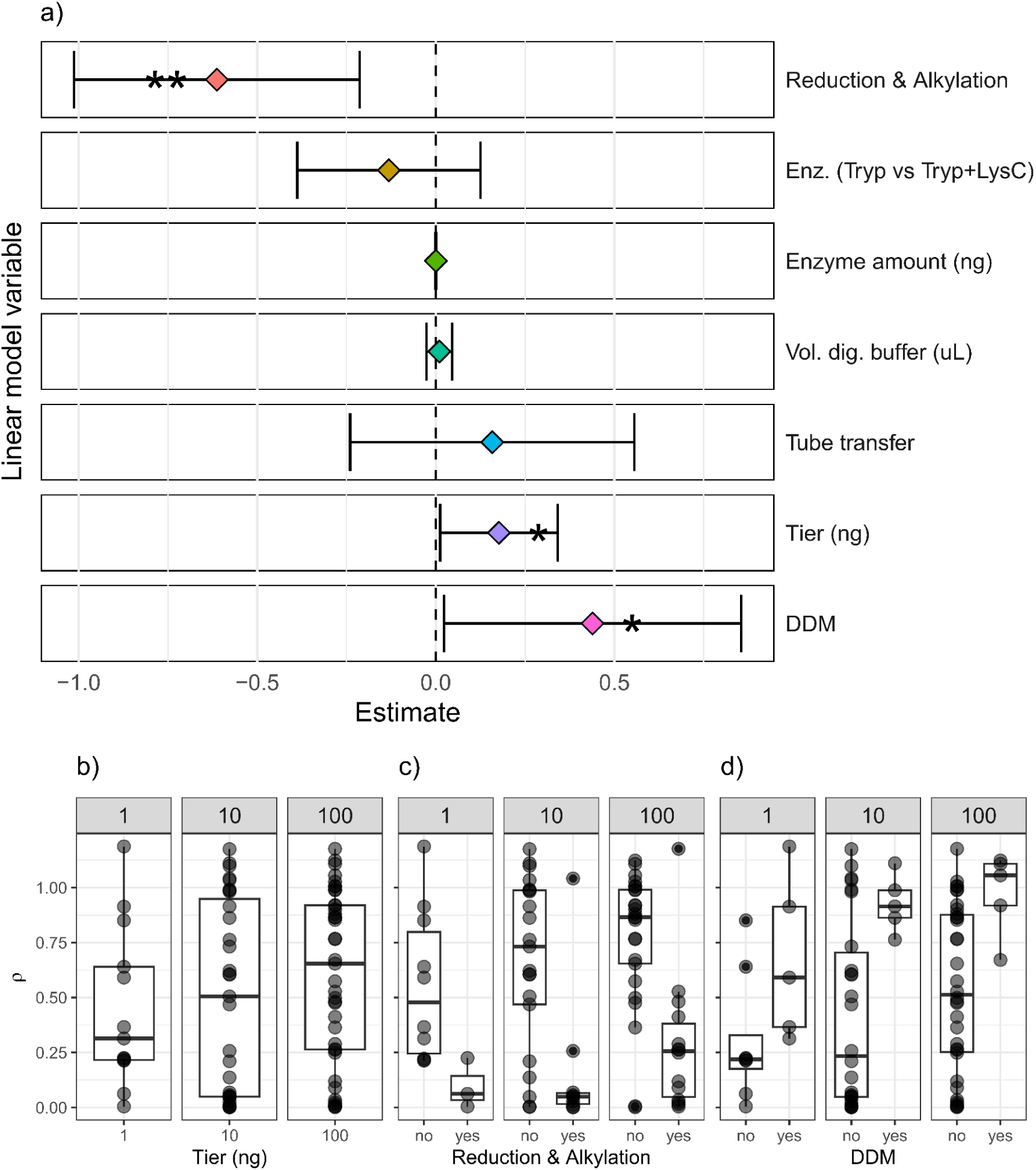
Linear model results and significant experimental variables. **a)** Graphical representation of the estimated coefficients from the linear model, with 95% confidence intervals for each explanatory variable used. Asterisks indicate statistically significant variables. Boxplots with *ρ* (y-axis) versus the starting sample amount (x-axis and grid columns) for each significant variable: **b)** tier in ng, **c)** reduction and alkylation, and **d)** DDM usage. (*ρ*: Recovery metric).

**Table 1.** Linear model summary. Coefficients of the linear model fitted on *ρ*_BC_, including estimates, standard errors, p-values, and 95% CIs (asterisks denote statistically significant variables). (*ρ*_BC_: Box-Cox transformed recovery; CI: Confidence intervals, Sign: Significance).

| Variable | Estimate | Standard error | t value | p value | Lower CI (2.5%) | Upper CI (97.5%) | Sign. |
| --- | --- | --- | --- | --- | --- | --- | --- |
| Reduction & Alkylation | -0.613 | 0.2003 | -3.061 | 0.003 | -1.0129 | -0.21 | ** |
| Enz. (Tryp vs. Tryp+LysC) | -0.131 | 0.1285 | -1.021 | 0.311 | -0.3879 | 0.125 |  |
| Enzyme amount (ng) | 0.0003 | 0.0005 | 0.4983 | 0.62 | -0.0008 | 0.001 |  |
| Vol. dig. buffer (μl) | 0.0099 | 0.0179 | 0.5516 | 0.583 | -0.0259 | 0.046 | |
| Tube transfer | 0.1579 | 0.1993 | 0.7924 | 0.431 | -0.24 | 0.556 |  |
| Tier (ng) | 0.1768 | 0.0825 | 2.1429 | 0.036 | 0.0121 | 0.341 | * |
| DDM | 0.4391 | 0.2084 | 2.1069 | 0.039 | 0.023 | 0.855 | * |

Overall, our inferential analysis validates previous recommendations to minimize sample manipulation steps, such as reduction and alkylation, and to incorporate DDM when working with low input amounts (Budnik et al., 2018; Nie et al., 2022). Importantly, these effects were consistently observed across laboratories, indicating that they represent robust and transferable principles rather than dataset specific observations. More broadly, we demonstrate that the proposed recovery framework provides a general strategy to enable fair inter-laboratory comparisons of sample preparation and can be readily extended to other multicentric studies.

## Conclusions

In the present study, we introduce ρ as a generalizable analytical framework to decouple protocol driven effects from analytical variability in multicentric studies.

We applied this conceptual framework to the analysis of PME13, demonstrating the critical role of sample preparation variables in HSP workflows, particularly at low protein inputs. Our results consistently show that the use of DDM enhances recovery, supporting its strong recommendation in HSP workflows. In contrast, reduction and alkylation and additional clean-up steps, although widely used, were associated with reduced performance, particularly at low input levels, likely due to increased sample loss. These findings further support the growing consensus that simplified, minimally handled workflows are optimal for maximizing sensitivity. Of note, while DIA generally outperforms DDA, according to our data, this advantage seems to diminish at low sample inputs (particularly regarding CV quality and the proportion of labs identifying *E. coli* proteins). This is likely because lower sample complexity reduces the inherent benefit of DIA over DDA. Finally, our study also highlights the importance of rigorous experimental practices, which should include blank experimental samples, to carefully control and monitor contamination when working at low input levels. In this context, the use of *E. coli* proved particularly valuable, as it enables both benchmarking of protocol performance and assessment of contaminant contributions across datasets.

Nevertheless, some limitations should be considered. A subset of datasets exhibited *ρ* values greater than one, likely reflecting technical issues such as suboptimal peptide resuspension in the Ref sample, which required the exclusion of these cases from downstream modeling. Additionally, the linear model was designed to identify global trends rather than to provide precise predictive power, and residual variability indicates that not all sources of variation are captured by the recorded metadata. Indeed, the metadata necessarily reflects only a subset of the numerous experimental factors that can influence performance in complex proteomics workflows.

Beyond the specific findings, this study establishes a general strategy for analyzing multicentric datasets, where normalization based metrics such as *ρ* can be used to minimize analytical confounding variables and enable meaningful cross laboratory sample preparation comparisons. This reference-based strategy may provide a useful foundation for benchmarking and standardization in other heterogeneous analytical workflows.

## Acknowledgements

The authors thank the EuPA for the continuous support to the Standardization Initiative. We would like to thank the whole ITS staff from IRB Barcelona for deploying and maintaining the file sharing and computation infrastructure (NextCloud, Open OnDemand, DIA-NN v.1.9.2) required to complete the present work.

IRB Barcelona Mass Spectrometry and Proteomics Core Facility is granted in the framework of the 2014-2020 ERDF Operational Programme in Catalonia, co-financed by the European Regional Development Fund (ERDF). Reference: IU16-015983.

The work was funded in part by grants to O.B.-S. from the Région Occitanie with European funds (Fonds Européens de Développement Régional, FEDER, REACT-EU program) and to O.B.-S. and C.C. from the French Ministry of Research with the Investissement d’Avenir Infrastructures Nationales en Biologie et Santé program (ProFI, Proteomics French Infrastructure project, ANR-10-INBS-08 and ANR-24-INBS-0015).

ProGenTomics acknowledges funding from the Research Foundation Flanders (FWO) [1278023N, I00882N] and Ghent University Special Research Fund [BOF21/DOC/076]. PI23CIII/00027 and PID2022-140307OB-I00 grants from the AES-ISCIII program and MCIN/AEI/10.13039/501100011033 cofounded by “ERDF A way of making Europe” funds, respectively to R.B. are also acknowledged.

This work was also supported by a grant of the Romanian Ministry of Research, Innovation and Digitization, CNCS - UEFISCDI, project numbers PN-IV-P8-8.3-ROMD-2023-0100 and PN-IV-P2-2.1-TE-2023-2082, within PNCDI IV.

ISPA’s Proteomics Unit has been funded by the Instituto de Salud Carlos III (ISCIII) (project IFCS22/00006) within the framework of the Mechanism for Recovery and Resilience (MRR) of the Next Generation EU funds.

This work was additionally supported by the competitive grants from the Spanish Ministry of Science, Innovation and Universities PLEC2022-009298, PLEC2022-009235, ”la Caixa” Foundation under the project code LCF/PR/HR22/52420019 and Comunidad de Madrid (IMMUNO-VAR, P2022/BMD-7333).

The CNB was supported by Grant CEX2023-001386-S funded by MICIU/AEI/ 10.13039/501100011033. Comunidad de Madrid Grants B2017/BMD-3817 and 2022/BMD-7232. MICIN PID2021-127496NB-100.

