## Supplementary material for "A generalizable normalization framework to decouple protocol and instrument effects: Application to high-sensitivity proteomics multicentric study (PME13)": Document1

### PME13
### EuPA standardization initiative: High sensitivity proteomics

#### Introduction

High sensitivity proteomics (HSP) has emerged in the last few years as a powerful tool in the field. Nowadays, mass spectrometry (MS) is sensitive enough to identify proteins at the single-cell level. On the other hand, MS sensitivity is approximately at attomole level ($10^{-18}$ moles), which involves hundred thousand ions. Since the median of protein copies in mammalian cells is around 18k copies per cell. MS is sensitive enough to analyze single cell samples. Indeed, most advanced studies provide about 1000 protein identifications per cell. In this journey towards HSP, sample preparation remains as one of the most crucial steps. Handling a low amount of material and transferring it into a mass spectrometer is still a challenge in proteomics [1].

Within the EuPA standardization initiative, we are launching a Proteomics Multicentric Experiment (PME13) focused on HSP. This multicentric study will provide us a unique opportunity to assess the performance and reproducibility of HSP experiments (from sample preparation to data analysis) within a specific lab and across multiple labs, platforms and sample preparation methodologies.

The reference samples to be analyzed in the study consist of a serial dilution of *E.coli* protein extract and peptide digest from 100 to 1 ng. The use of standard *E.coli* samples will allow us not only to monitor the number of identified proteins from *E.coli* but also to track potential human protein contaminants during sample preparation, which is one of the drawbacks/pitfalls of the whole procedure. The final aim of the study is to test the performance and reproducibility of HSP protocols and assess the usefulness of the *E.coli* standard to set up and troubleshoot HSP workflows.

#### Objectives

The main goal of PME13 is to benchmark protocols for HSP analysis and to provide tools to facilitate HSP implementation and to assess laboratory performance on HSP analyses. Specifically:

1. To assess the robustness and reproducibility of different HSP protocols.
2. To deliver references to assess performance on HSP analysis.
3. To provide a reference sample for HSP protocol standardization.
4. To provide a data analysis workflow.

#### PME13 study

##### Sample

There will be two sample sets: ***protein*** and ***peptide***.

- **Protein (AKA HS sample):** *E.coli* protein extract (BioRad: *E.coli* Protein Sample -- 163-2110) at different amounts: 100, 10, 1 ng. For each protein concentration 4 replicates will be provided (4x3 = 12 samples)

- **Peptide (AKA Ref sample):** Tryptic digest of *E.coli* protein extract (BioRad: *E.coli* Protein Sample -- 163-2110) at different amounts: 100, 10, 1 ng. For each peptide concentration 4 replicates will be provided (4x3 = 12 samples).

Participants will process protein samples and analyze the 24 samples provided (protein and peptide) by LCMS. You should run the samples by data dependent acquisition (DDA) but, optionally, you can also analyze the samples by data independent acquisition (DIA). If you choose the second option you will receive 48 samples instead of 24.

#### Data analysis

All database searches and data analysis results will be centralized. You will receive an email with a NextCloud link to upload your MS files. Each participant will have a distinct link that points to a separated NextCloud folder (this system should minimize the chances of Spectrum File scrambling).

Database searches (using MaxQuant v2.3.X.X [2] or DIA-NN v1.8.X [3], later upgraded to v1.9.2) and the subsequent data analysis will be performed at IRB Barcelona. We will put the focus on the number of *E.coli* and human proteins identified (human protein identifications will be considered contaminants). We will also analyze the reproducibility among replicates within each sample and laboratory. Moreover, we will compare results from same-injected amount samples coming from peptide or protein samples. Finally, we will compare the performance drop as we reduce the amount of starting material in protein and peptide samples. Peptide samples will serve as internal lab reference and will be compared with protein samples to calculate a digestion protein recovery. This recovery will be used to compare lab sample preparation performance.

These analyses will help us to determine which are the key factors for an optimal sample preparation. We will extract final conclusions on the best sample preparation protocols and LCMS platforms according to the approaches used by each participating lab.

#### Outcome

A workshop will be organized to share and discuss PME13 results once the study is concluded. A preliminary analysis will be presented at ProteoAix 2023 (June 20-23) and the first draft of the results will be discussed at BSPR-EuPA 2023 (July 17-20).

The final goal would be to prepare a manuscript in which all participants will appear as authors.

### How to participate in the PME13 study?

#### Phase 0: Fill the request form

If you want to join the PME13 study you should fill the following enrollment form (closed on May 10th 2023):

https://forms.gle/qXJp646gXZRe1QYUA

After filling the form, you will receive an email from with your **Lab ID**, your NextCloud **uploading link** and a **metadata spreadsheet**. Each participant will have a distinct link that points to a separated NextCloud folder. As PME13 pretends to be an anonymized study, you should use the Lab ID received  to name your LCMS files. Finally, the metadata spreadsheet will enable us to correlate the experiment performance with the sample preparation information.

#### Phase 1: Sample shipping and naming

Samples will be shipped to your lab during April 2023. You will receive 24 PCR tubes with all the samples and replicates. You will digest the 12 protein samples and analyze all the 24 samples. If you choose to also perform the experiment by DIA, you will process 24 additional samples (48 in total).

Sample labeling and naming is crucial to facilitate the centralized analysis. For this reason, we will kindly ask you to name your samples as shown in the tables below.

| Lab ID | Sample type | Acquisition method | Peptide or protein amount (ng) | Replicate | LCMS file name | Injection order |
|---|---|---|---|---|---|---|
| Lab00 | Prot | DDA | 100 | 1 | Lab00_Prot_DDA_100_R1 | 17 |
| Lab00 | Prot | DDA | 100 | 2 | Lab00_Prot_DDA_100_R2 | 18 |
| Lab00 | Prot | DDA | 100 | 3 | Lab00_Prot_DDA_100_R3 | 19 |
| Lab00 | Prot | DDA | 100 | 4 | Lab00_Prot_DDA_100_R4 | 20 |
| Lab00 | Prot | DDA | 10 | 1 | Lab00_Prot_DDA_10_R1 | 9 |
| Lab00 | Prot | DDA | 10 | 2 | Lab00_Prot_DDA_10_R2 | 10 |
| Lab00 | Prot | DDA | 10 | 3 | Lab00_Prot_DDA_10_R3 | 11 |
| Lab00 | Prot | DDA | 10 | 4 | Lab00_Prot_DDA_10_R4 | 12 |
| Lab00 | Prot | DDA | 1 | 1 | Lab00_Prot_DDA_1_R1 | 1 |
| Lab00 | Prot | DDA | 1 | 2 | Lab00_Prot_DDA_1_R2 | 2 |
| Lab00 | Prot | DDA | 1 | 3 | Lab00_Prot_DDA_1_R3 | 3 |
| Lab00 | Prot | DDA | 1 | 4 | Lab00_Prot_DDA_1_R4 | 4 |
| Lab00 | Pep | DDA | 100 | 1 | Lab00_Pep_DDA_100_R1 | 21 |
| Lab00 | Pep | DDA | 100 | 2 | Lab00_Pep_DDA_100_R2 | 22 |
| Lab00 | Pep | DDA | 100 | 3 | Lab00_Pep_DDA_100_R3 | 23 |
| Lab00 | Pep | DDA | 100 | 4 | Lab00_Pep_DDA_100_R4 | 24 |
| Lab00 | Pep | DDA | 10 | 1 | Lab00_Pep_DDA_10_R1 | 13 |
| Lab00 | Pep | DDA | 10 | 2 | Lab00_Pep_DDA_10_R2 | 14 |
| Lab00 | Pep | DDA | 10 | 3 | Lab00_Pep_DDA_10_R3 | 15 |
| Lab00 | Pep | DDA | 10 | 4 | Lab00_Pep_DDA_10_R4 | 16 |
| Lab00 | Pep | DDA | 1 | 1 | Lab00_Pep_DDA_1_R1 | 5 |
| Lab00 | Pep | DDA | 1 | 2 | Lab00_Pep_DDA_1_R2 | 6 |
| Lab00 | Pep | DDA | 1 | 3 | Lab00_Pep_DDA_1_R3 | 7 |
| Lab00 | Pep | DDA | 1 | 4 | Lab00_Pep_DDA_1_R4 | 8 |

Prot = Protein, Pep = Peptide. If the "Acquisition method" is DIA instead of DDA, the entries in the "LCMS file name" column should be changed accordingly (i.e. "_DIA_" instead of "_DDA_").

#### Phase 2: Sample preparation

You will digest the 12 protein samples. We recommend adapting the standard methods considering you will work with low protein amounts. When possible, you should avoid tube transferring. Please, use trypsin or LysC/trypsin as a digestion enzyme. If you don't have a robot to prepare samples with low amounts, a good point to start is to adapt the first SCoPE sample preparation protocol to PME13 samples (without the TMT labeling part).

You will resuspend the 12 peptide samples. We recommend 0.1 % formic acid.

#### Phase 3: Sample LCMS analysis

You will analyze in your LCMS system the 24 samples. You will inject all the amount of sample for the aliquots 100, 10 and 1 ng. ***Samples should be injected in increasing order of concentration as shown in the table above: first 1 ng, then 10 ng and finally 100 ng samples.***

You can adapt your preferred LC and MS parameters for high sensitivity experiments to each one of the starting amounts tiers: 1 ng, 10 ng or 100 ng. For example, you can choose your LC gradient and buffers, the MS acquisition parameters at the same tier, but ***you must use the same LCMS setup for all the replicates comprising the same starting amount (note the different colors in the table above).*** Please, name the LCMS files as described in the table above.

Choosing DDA as acquisition method is mandatory. Optionally, you can also acquire the extra 24 samples by DIA if you wish.

#### Phase 4: Data analysis

The data analysis will be centralized at the MSPCF from IRB Barcelona. In order to enable them to complete the data analysis, you should share with them the raw files generated by your LC/MS system, as well as some additional metadata regarding your experimental approach. Feel free to contact them if you have any issues along the way.

### Sharing the Spectrum Files

Please follow the instructions below to share your Spectrum Files:

1. Create a single zip file containing all your Spectrum Files. This process might take a substantial amount of time depending on the total size of the files and the particular zipping tool (please be patient). **Note**: If you are using an Astral, or any other instrument generating huge spectrum files, skip the zipping step.
2. Once the zipping process is finished, follow the NextCloud link. A window like this should appear in your web browser:

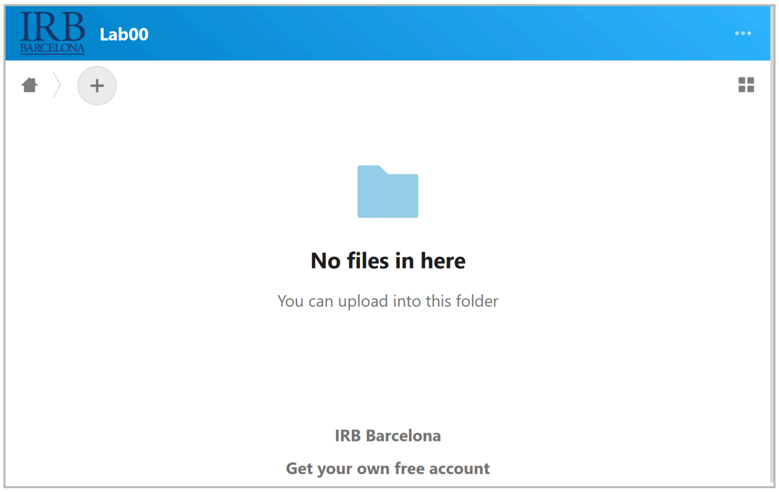


3. Check if your Lab ID matches the Lab ID label appearing on the top left corner of the window (notice the “Lab00” label on the picture above). Contact us if your Lab ID doesn’t match the Lab ID label from NextCloud.
4. Drag-and-drop your single zip file containing all your Spectrum Files into the NextCloud window:

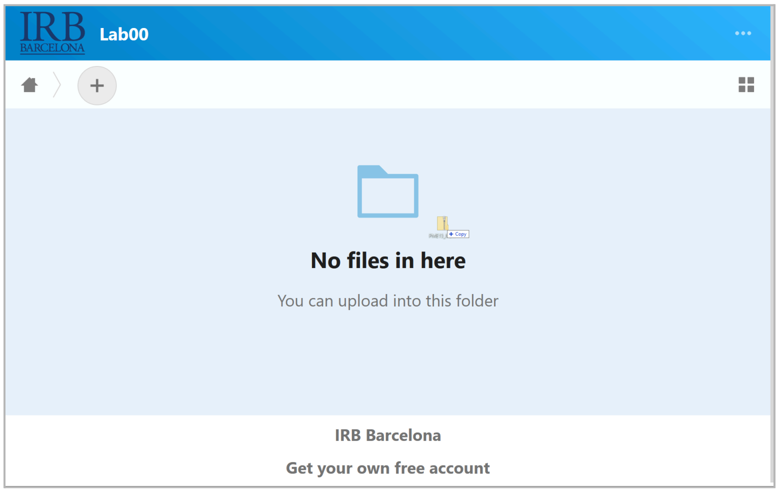


5. Wait until the uploading process is completed (notice the progression bar on the top left corner). This process might take a substantial period of time depending on the total size of the files and your network bandwidth (please be patient)

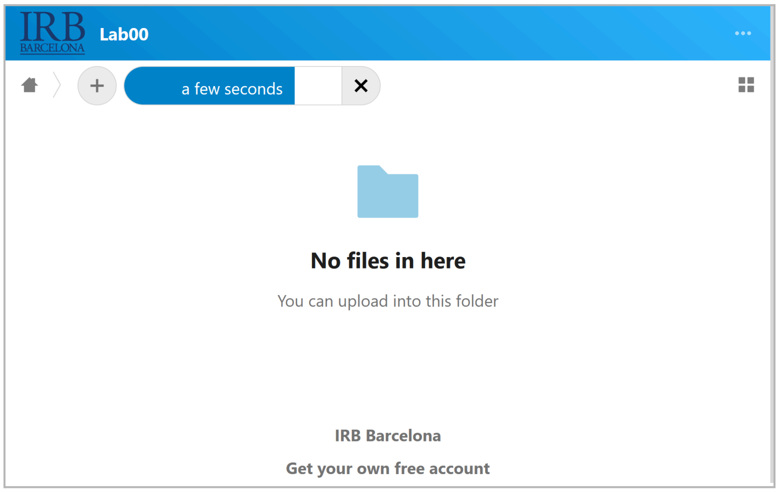


6. Check that your single zip file containing all your Spectrum Files is successfully uploaded (notice the file should appear listed):

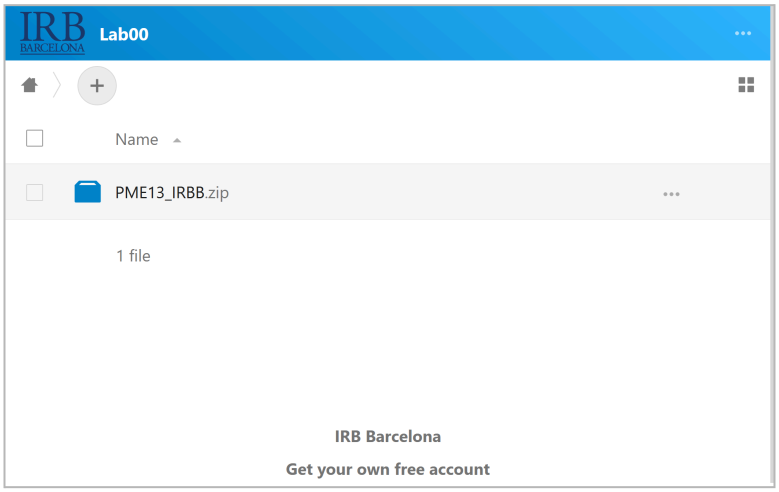


#### Sharing the Metadata

In order to correlate the results with the sample preparation information, we ask you to carefully fill the metadata spreadsheet you will receive at the time you join the PME13. In this spreadsheet, you will find a text box with some instructions. In any case, if you have any problem when filling the metadata spreadsheet, you can directly contact us. Please, follow the steps (starting from step 2) described in subsection "*Phase 4: Data analysis (Sharing the Spectrum Files)*" to upload your metadata spreadsheet to NextCloud. Feel free to also share with us a text file describing your particular sample preparation protocol.

#### Phase 5: Final meeting with all the participants

Once we have collected and analyzed all the data a meeting will be proposed to show and discuss the results. A preliminary analysis will be presented at ProteoAix 2023 (June 20-23) and the first draft of the results will be discussed at BSPR-EuPA 2023 (July 17-20).

#### Due dates and deadlines

| | |
|---|---|
| **PME13 announcement and launching** | 15/03/2023 |
| **Enrollment form (Participants)** | 15/03/2023 - 30/04/2023 |
| **Sample shipping (Organizers)** | 11/04/2023 - 30/04/2023 |
| **Upload spectrum files and Metadata form (Participants)** | 15/06/2023 |
| **Data analysis (Organizers)** | June 2023 |
| **Preliminary analysis** | ProteoAix 2023 |
| **First draft** | BSPR-EuPA 2023 |

#### FAQs

**Q**: How long will it take for the samples to arrive?
**A**: It is difficult to estimate because it will not depend exclusively on us (courier service, geographical distance to the destination, etc). We will send the samples as the registrations arrive (the sooner you fill out the registration form, the sooner you will receive the samples).

**Q**: Can I run the experiment using DIA only?
**A**: No. There are just two options: The first one: DDA (mandatory) and the second one: DDA + DIA (optional).

**Q**: The BioRad reference (163-2110) for the E.coli peptide sample is the same as for the E.coli protein sample. There is some mistake in those references?
**A**: No. For the protein sample we will use *E.coli* protein extract from BioRad as it is, and for the peptide sample we will start from this same *E.coli* protein extract and we will perform its tryptic digestion. In any case, you do not have to worry about any of this because we will take care of sending you both samples already prepared.

**Q**: I filled the enrollment form and I did not receive any email. Is this normal?
**A**: Yes. Before sending the welcoming email, we check if the participant data is trustworthy. Therefore, it is normal to wait a few days before receiving the welcoming email.

**Q**: I just received the samples. Should I notify somebody?

**Q**: I uploaded the zip file with the spectrum files via NextCloud (and/or the metadata spreadsheet) but they don't appear listed as depicted in this document (*Phase 4: Data analysis; Sharing the Spectrum Files; Step number 6*). Is this normal?
**A**: Yes it is. Some users have reported that the uploaded files are not visible exactly as shown in this document. However, a message like: "*Files uploaded: PME13_IRBB.zip*" should appear instead, and it is a successful uploading process confirmation message.
