## Supplementary Figures for "A generalizable normalization framework to decouple protocol and instrument effects: Application to high-sensitivity proteomics multicentric study (PME13)"


Figure S1
Map of participating countries.
Figure S2
- a) Contaminant presence breakdown for data dependent acquisition (DDA) at the peptide spectrum match level (PSM).
- b) Contaminant presence breakdown for data independent acquisition (DIA) at the precursor level.

Figure S3
- a) Coefficients of variation breakdown for data dependent acquisition (DDA) at the peptide spectrum match level (PSM).
- b) Coefficients of variation breakdown for data independent acquisition (DIA) at the precursor level.

Figure S4
- a) Outcome metrics analysis breakdown for data dependent acquisition (DDA).
  - i) Exploris 240
  - ii) Exploris 480
  - iii) LTQ Orbitrap Pro
  - iv) Orbitrap Astral
  - v) Orbitrap Eclipse
  - vi) Q Exactive
  - vii) timsTOF-FLEX
  - viii) timsTOF-SCP
  - ix) ZenoTOF 7600
- b) Outcome metrics analysis breakdown for data independent acquisition (DIA).
  - i) Exploris 480
  - ii) Orbitrap Astral
  - iii) Orbitrap Eclipse
  - iv) timsTOF-FLEX
  - v) timsTOF-SCP
  - vi) ZenoTOF 7600

**Figure S1**

**Map of participating countries.**

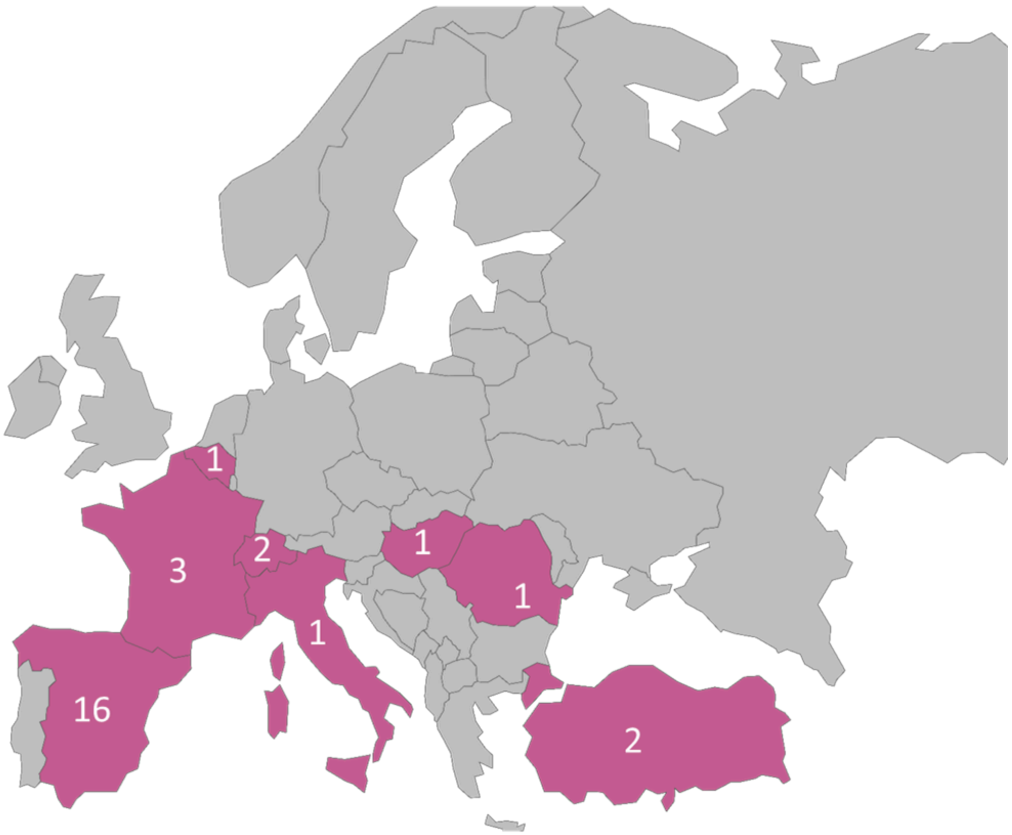


**Figure S1.** Europe map showing the countries involved in the Proteomic Multicentric Experiment 13 (PME13) in purple. Numbers indicate the participation per country.

### Figure S2

**a) Contaminant presence breakdown for data dependent acquisition (DDA) at the peptide spectrum match level (PSM).**

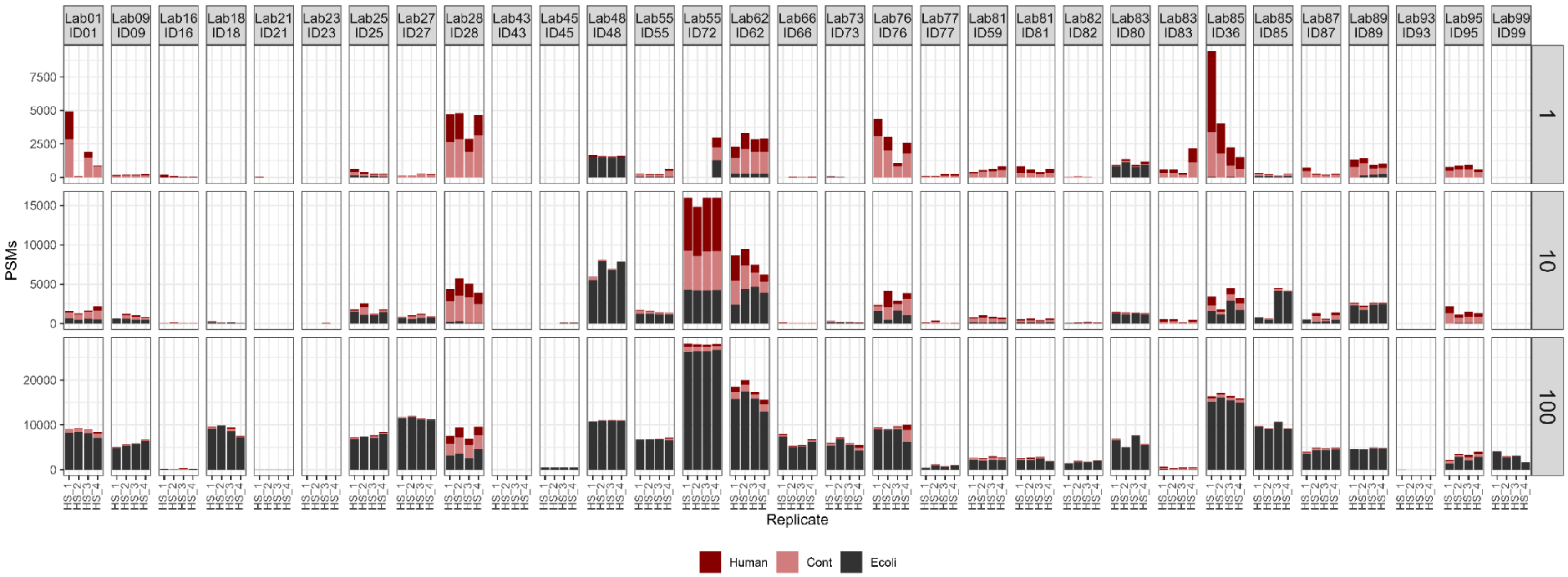


**Figure S2a**. Stacked bar plots with the number of PSMs (y-axis) identified across the four replicates of the High Sensitivity (HS) sample (x-axis), for each dataset (grid columns) and each tier in ng (grid rows). The color indicates the database.

**b) Contaminant presence breakdown for data independent acquisition (DIA) at the precursor level.**

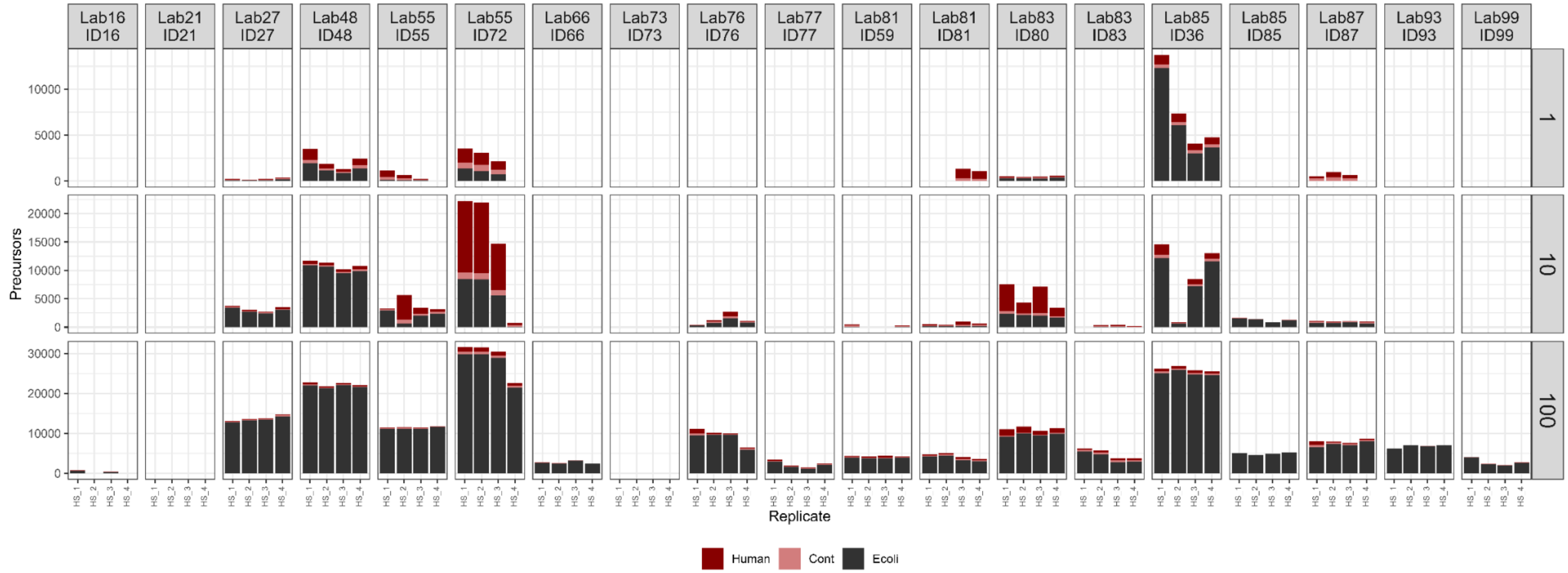


**Figure S2b**. Stacked bar plots with the number of precursors (y-axis) identified across the four replicates of the High Sensitivity (HS) sample (x-axis), for each dataset (grid columns) and each tier in ng (grid rows). The color indicates the database.

**Figure S3**

**a) Coefficients of variation breakdown for data dependent acquisition (DDA) at the peptide spectrum match level (PSM).**

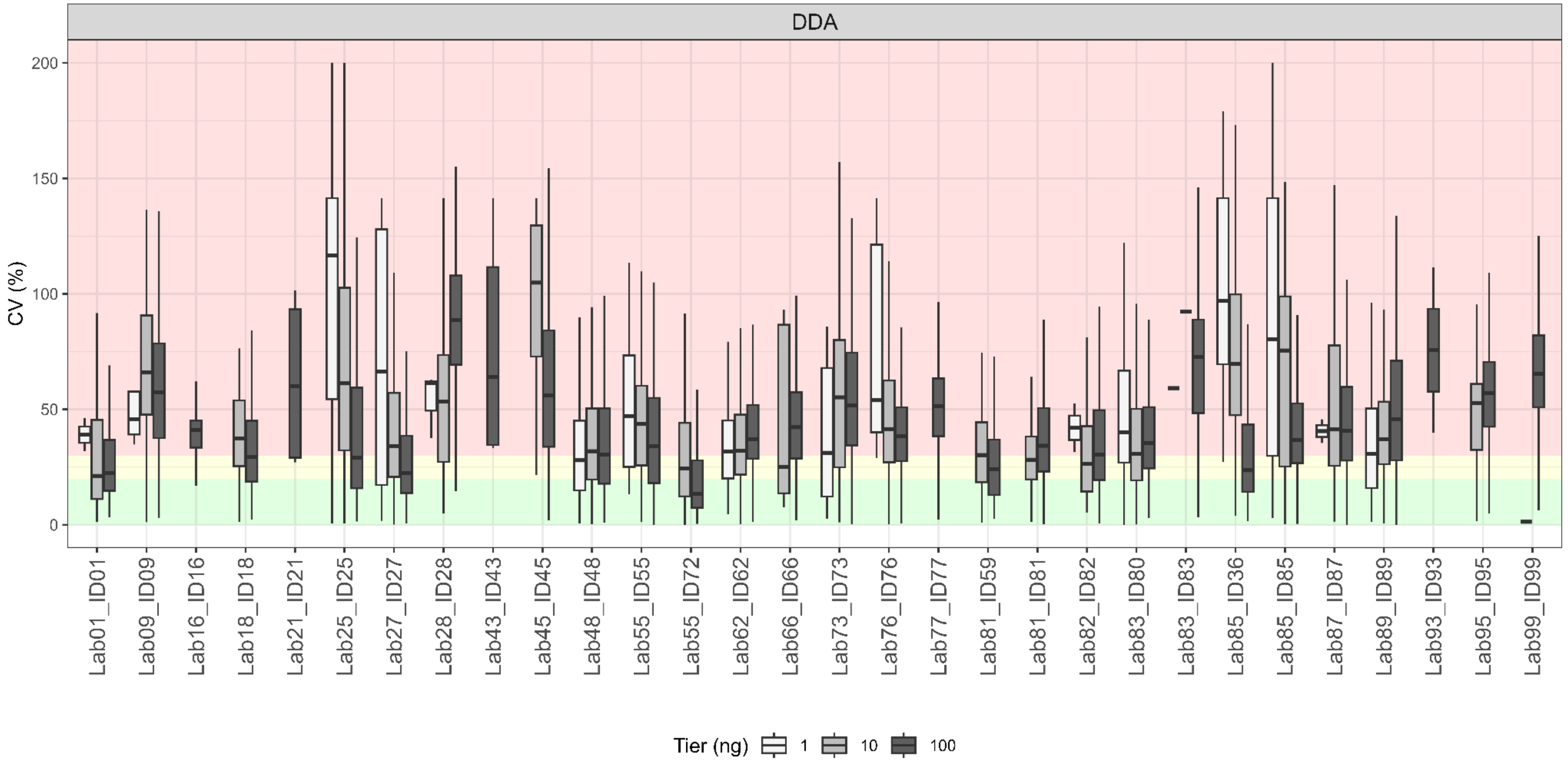


**Figure S3a**. Boxplots with coefficients of variation (CV), expressed as a percentage, for *E. coli* protein group (PG) identified (y-axis), across laboratories and datasets (x-axis). The gray shade differentiates the initial sample amount. The CVs shown in panels **a)** and **b)** correspond to data dependent and data independent acquisition modes (DDA and DIA, respectively). The background colors indicate CV ranges: green: CV < 20%, yellow: 20% < CV < 30%, red: CV > 30%.

**b) Coefficients of variation breakdown for data independent acquisition (DIA) at the precursor level.**

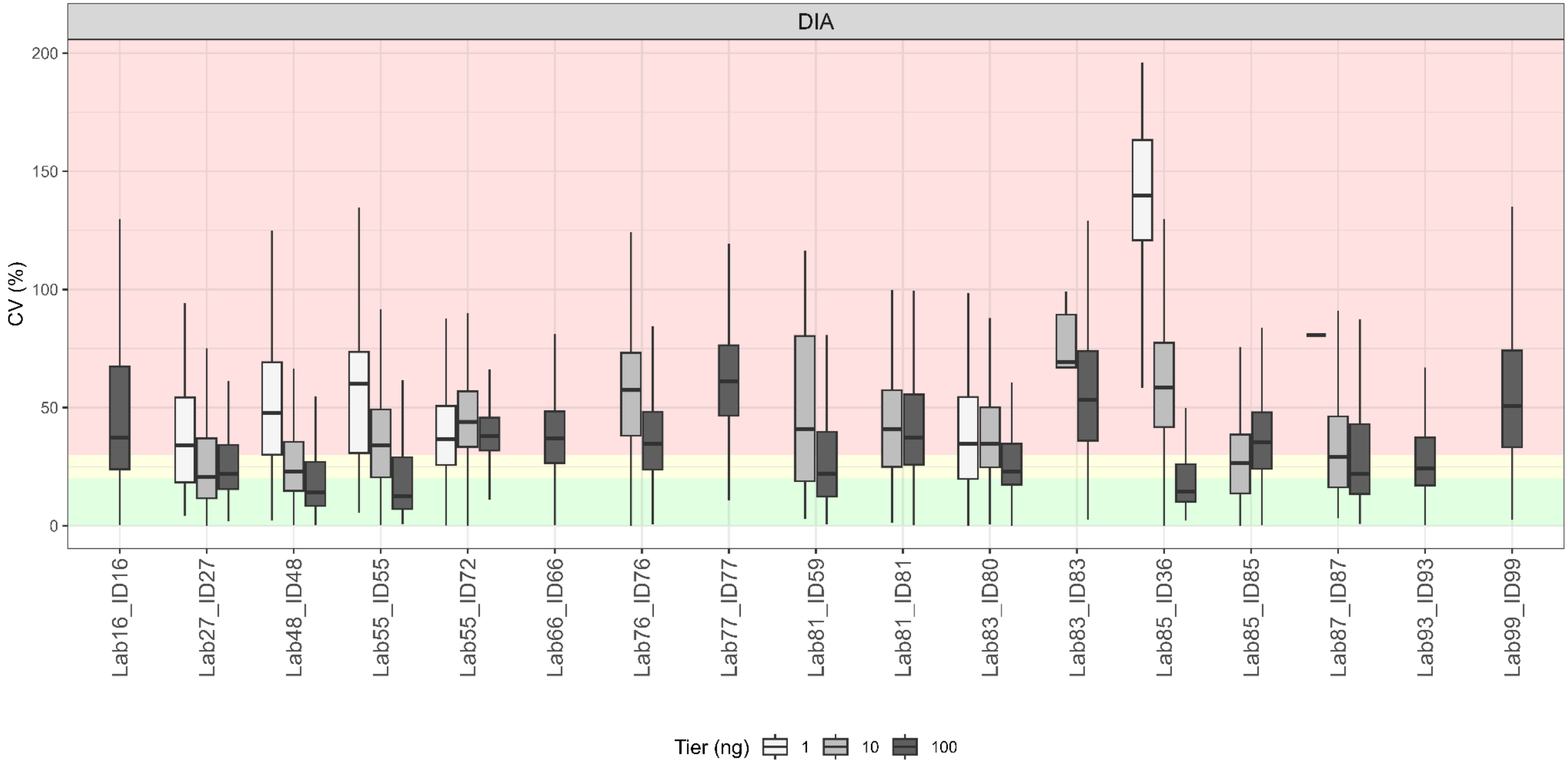


**Figure S3a**. Boxplots with coefficients of variation (CV), expressed as a percentage, for *E. coli* protein group (PG) identified (y-axis), across laboratories and datasets (x-axis). The gray shade differentiates the initial sample amount. The CVs shown in panels **a)** and **b)** correspond to data dependent and data independent acquisition modes (DDA and DIA, respectively). The background colors indicate CV ranges: green: CV < 20%, yellow: 20% < CV < 30%, red: CV > 30%.

**Figure S4**

**a) Outcome metrics analysis breakdown for data dependent acquisition (DDA).**

*i) Exploris 240*

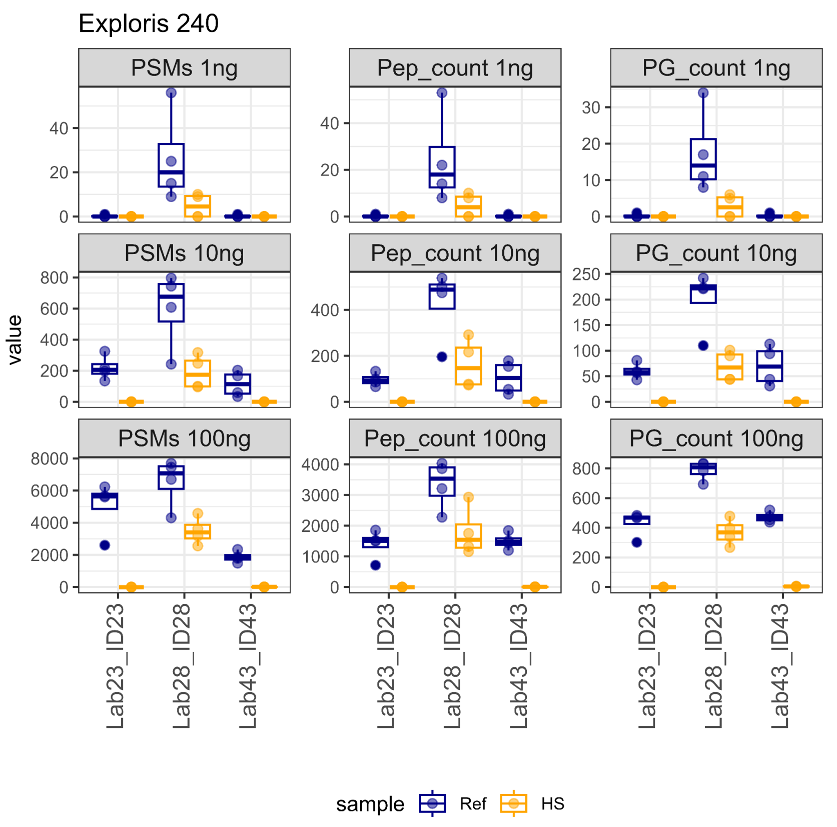


| Dataset | ID23 | ID28 | ID43 |
|---|---|---|---|
| Lab | Lab23 | Lab28 | Lab43 |
| Tube.transfer | 1 | 0 | 1 |
| Reduction | 1 | 1 | 1 |
| Reduction.reagent | DTT | TCEP 100mM(2Î¼l) | 10 mM TCEP; 75 Â°C, 15 min; 1 ul |
| Alkylation | 1 | 1 | 1 |
| Alkylation.Reagent | CHLOROACETAMI DE | CAA 200mM(2Î¼l) | 20 mM IAM; room temp., 30 min, dark; 1 ul |
| Enzyme | Tryp | Tryp | Tryp+LysC |
| Enzyme.amount..ng. | 1\|0.1\|0.01 | 500 | 2.5\|0.25\|0.025 |
| Digestion.Buffer | 0,9 M UREA IN 100 Mm TRIS-HACL PH 8,5, 1 Mm CaCl2 | TEAB 100mM | 20 mM ammonium bicarbonate |
| Volume.of.digestion.buffer..uL. | 450 | 30 | 10 |
| HPLC | NANO ULTIMATE 3000 | Ultimate 3000 | Thermo Scientific UltiMate 3000 nLC |
| Gradient..min. | 157 | 60 | 120 |
| Column length (cm) | 50 | 50 | 50 |
| MS2.detector | ORBITRAP | High | Orbitrap (high resolution) |
| MS_newName | Exploris 240 | Exploris 240 | Exploris 240 |

*ii) Exploris 480*

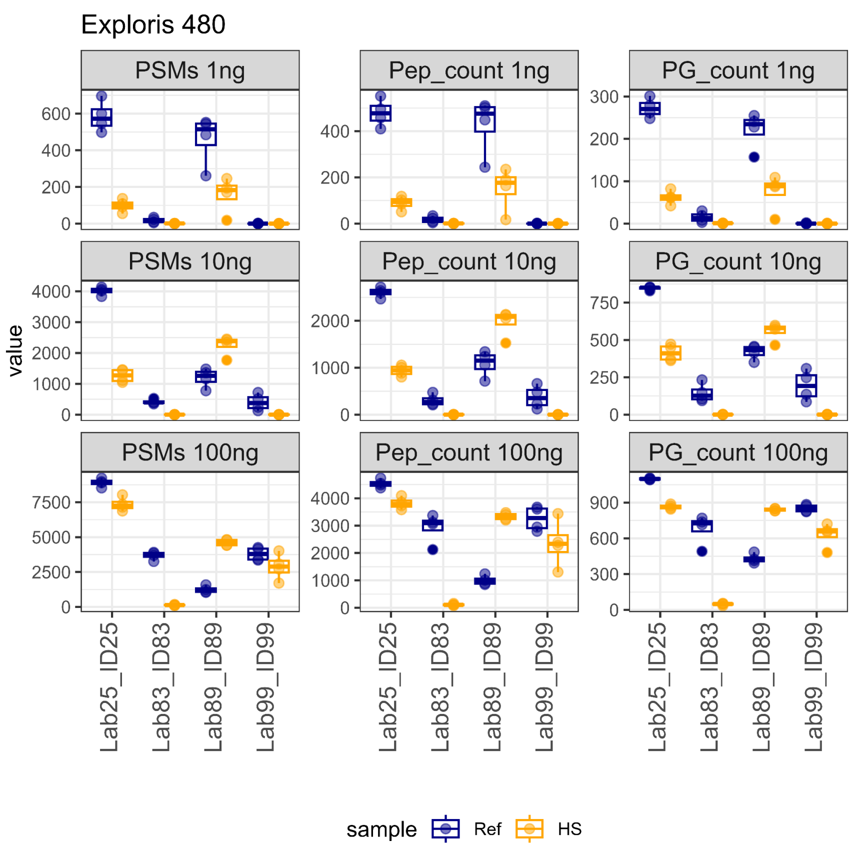


| Dataset | ID25 | ID83 | ID89 | ID99 |
|---|---|---|---|---|
| **Lab** | **Lab25** | **Lab83** | **Lab89** | **Lab99** |
| **Tube.transfer** | 0 | 0 | 0 | 0 |
| **Reduction** | 0 | 1 | 0 | 0 |
| **Reduction reagent** | NA | 4 ÂµL of 100 mM TCEP, 10 mM TCEP final concentration | NA | NA |
| **Alkylation** | 0 | 1 | 0 | 0 |
| **Alkylation** | NA | 4 ÂµL of 400 mM 2-chloroacetamide4 0 mM 2-chloroacetamide final concentration | NA | NA |
| **Reagent** | | | | |
| **Enzyme** | Tryp | Tryp+LysC | Tryp | Tryp |
| **Enzyme.amount..ng.** | 10 | 500 | 20 | 50\|5\|0.5 |
| **Digestion.Buffer** | TEABC 50 mM | 50 mM ABC pH 8.0 | ProteaseMax 0.05%, TEAB 100 mM | ABC 50mM |
| **Volume.of.digestion.buffer..uL.** | 1 | 20 | 10 | 20 |
| **HPLC** | RSLC U3000 (PepMap C18 75umx250mm) | Vanquish Neo UHPLC | EASY nLC-1200 | Ez100 |
| **Gradient..min.** | 122 | 76 | 65\|20\|20 | 15 |
| **Column length (cm)** | 25 | 50 | 25 | 25 |
| **MS2.detector** | High resolution (Orbitrap)120K (MS) 30K (MSÂ²) | | | |
| **MS_newName** | Exploris 480 | Exploris 480 | Exploris 480 | Exploris 480 |
| **FAIMS** | yes | no | yes | no |

*iii)* *LTQ Orbitrap Velos Pro*

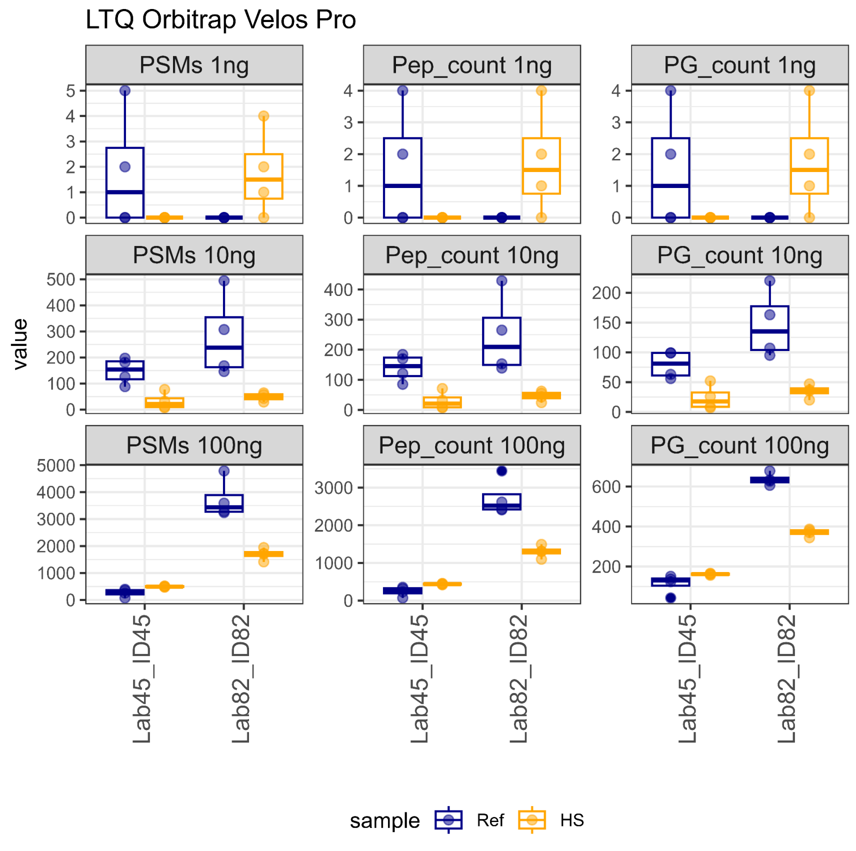


| Dataset | ID45 | ID82 |
|---|---|---|
| **Lab** | **Lab45** | **Lab82** |
| **Tube.transfer** | 0 | 0 |
| **Reduction** | 0 | 0 |
| **Reduction.reagent** | NA | NA |
| **Alkylation** | 0 | 0 |
| **Alkylation.Reagent** | NA | NA |
| **Enzyme** | Tryp+LysC | Tryp |
| **Enzyme.amount..ng.** | 4 | 25 |
| **Digestion.Buffer** | 10 mM ABC | Tris pH8 |
| **Volume.of.digestion.buffer..uL.** | 10 | 10 |
| **HPLC** | Easy-nanoLC II | Easy nLC-1200 |
| **Gradient..min.** | 90 | 120\|90\|60 |
| **Column length (cm)** | 15 | 25 |
| **MS2.detector** | LTQ | High |
| **Notes** | Used a 90-min 2-30% solv B. gradient with MS Orbitrap scan and LTQ detection of the ion fragments following CID. | |
| **MS_newName** | LTQ Orbitrap Velos Pro | LTQ Orbitrap Velos Pro |

*iv) Orbitrap Astral*

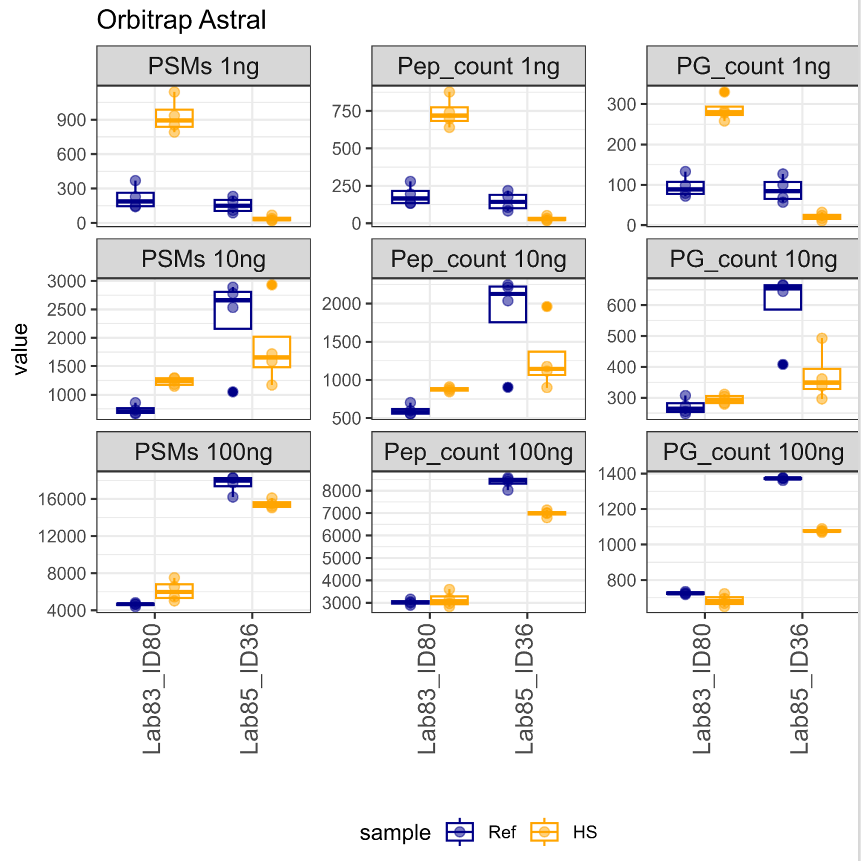


| Dataset | ID36 | ID80 |
|---|---|---|
| Lab | Lab36 | Lab80 |
| Tube.transfer | 1 | 0 |
| Reduction | 0 | 0 |
| Reduction.reagent | | |
| Alkylation | 0 | 0 |
| Alkylation.Reagent | | |
| Enzyme | Tryp | Tryp+LysC |
| Enzyme.amount..ng. | 50 | 50 |
| Digestion.Buffer | ABC 0.2M | TEAB 50mM, 0.01% DMM |
| Volume.of.digestion.buffer..uL. | 3 | 5 |
| HPLC | Vanquish Neo | Vanquish Neo |
| Gradient..min. | 30 | 18 |
| Column length (cm) | 50 | 5.5 |
| MS2.detector | Astral | Astral, 80000 |
| MS_newName | Orbitrap Astral | Orbitrap Astral |

*v)* *Orbitrap Eclipse*

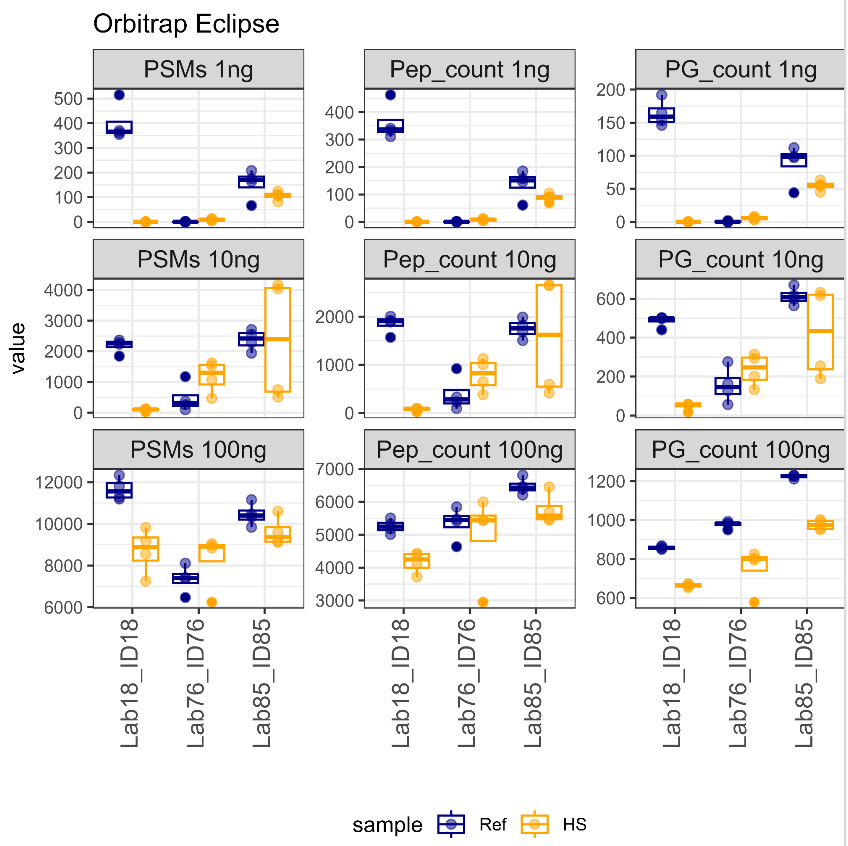


| Dataset | ID18 | ID76 | ID85 |
|---|---|---|---|
| Lab | Lab18 | Lab76 | Lab85 |
| Tube.transfer | 0 | 1 | 1 |
| Reduction | 0 | 0 | 0 |
| Reduction.reagent | NA | NA | NA |
| Alkylation | 0 | 0 | 0 |
| Alkylation.Reagent | NA | NA | NA |
| Enzyme | Tryp+LysC | Tryp | Tryp |
| Enzyme.amount..ng. | 10 | 75 | 50 |
| Digestion.Buffer | 7.5% ACN, 100 mM Ammonium bicarbonate | 100mM Ammonium bicarbonate | ABC 0.2M |
| Volume.of.digestion.buffer..uL. | 20 | 22 | 3 |
| HPLC | Evosep | Evosep | Thermo nLC-1200 |
| Gradient..min. | 31 | 44 | 60 |
| Column length (cm) | 15 | 8 | 50 |
| MS2.detector | IT | Low Res. | ION TRAP |
| Notes | | | Inject 90% sample |
| MS_newName | Orbitrap Eclipse | Orbitrap Eclipse | Orbitrap Eclipse |

*vi)* *Q Exactive*

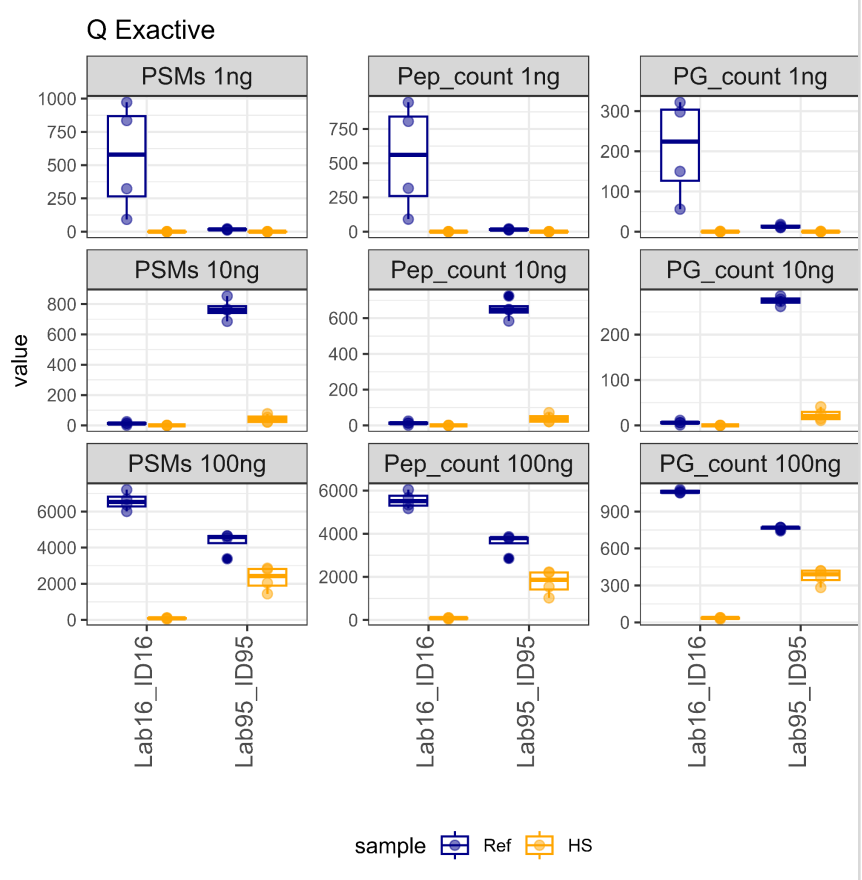


| Dataset | ID16 | ID95 |
|---|---|---|
| Lab | Lab16 | Lab95 |
| Tube.transfer | 0 | 0 |
| Reduction | 1 | 1 |
| Reduction.reagent | DTT,10mM final, 2ul | TCEP 100mM, 2ÂµL, vol.final 40ÂµL |
| Alkylation | 1 | 1 |
| Alkylation.Reagent | CAA,20mM final, 4,4ul | CAA 200mM, 2ÂµL, vol.final 40ÂµL |
| Enzyme | Tryp | Tryp |
| Enzyme.amount..ng. | 5\|0.5\|0.05 | 500 |
| Digestion.Buffer | 50 mM ABC | TEAB 100 mM |
| Volume.of.digestion.buffer..uL. | 158.4 | 20 |
| HPLC | Dionex Ultimate 3000 NCS-3500RS | Easy-nLC 1000 |
| Gradient..min. | 90\|90\|40 | 60 |
| Column length (cm) | 40 | 50 |
| MS2.detector | High (70,000) | High (70K) |
| Notes | samples dissolved in 20ul 8M urea in Tris-Cl upon arrival, desalted using C18 StageTips after digest | Samples were not cleaned-up |
| MS_newName | Q Exactive | Q Exactive |

*vii) timsTOF-FLEX*

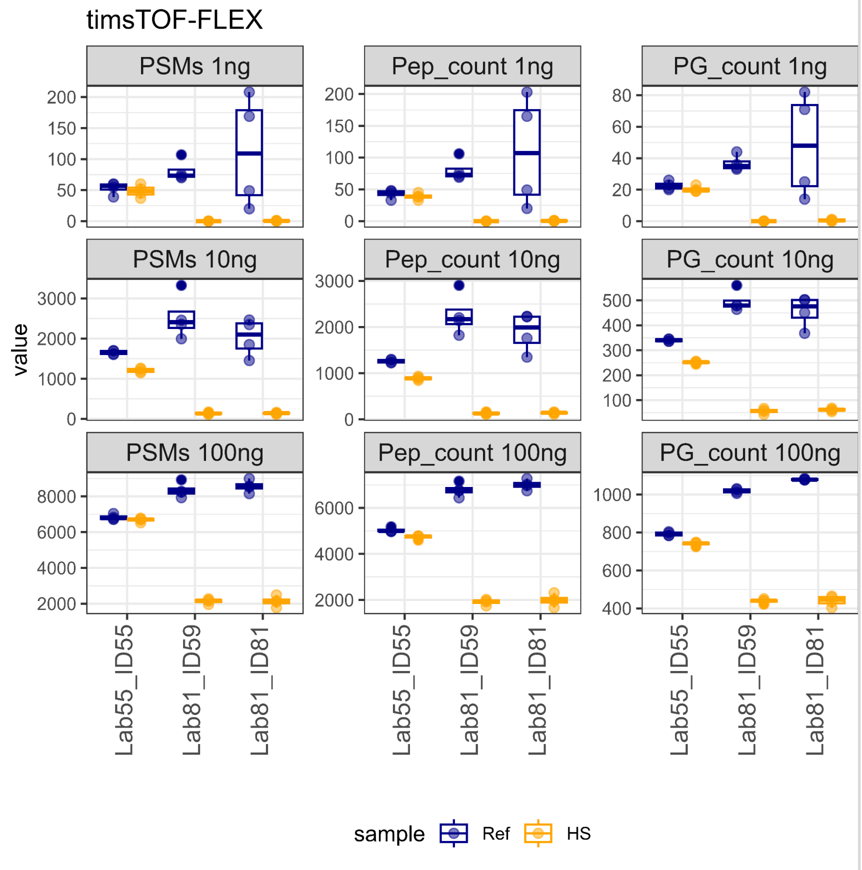


| Dataset | ID55 | ID59 | ID81 |
|---|---|---|---|
| Lab | Lab55 | Lab81 | Lab81 |
| Tube.transfer | 0 | 1 | 1 |
| Reduction | 0 | 1 | 1 |
| Reduction.reagent | NA | iSTkit | sp3_iSTkit |
| Alkylation | 0 | 1 | 1 |
| Alkylation.Reagent | NA | iSTkit | sp3_iSTkit |
| Enzyme | Tryp+LysC | Tryp+LysC | Tryp+LysC |
| Enzyme.amount..ng. | 5 | NA | NA |
| Digestion.Buffer | 50 mM ABC | iSTkit | sp3_iSTkit |
| Volume.of.digestion.buffer..uL. | 18 | | |
| HPLC | EVOSEP ONE | Evosep | Evosep |
| Gradient..min. | 21 | 21 | 21 |
| Column length (cm) | 8 | 8 | 8 |
| MS2.detector | HIGH RESOLUTION | HIGH | HIGH |
| Notes | | iST kits from Preomics to perform the digestions. | sp3 add-on to iST kits from Preomics |
| MS_newName | timsTOF-FLEX | timsTOF-FLEX | timsTOF-FLEX |

*viii) timsTOF-SCP*

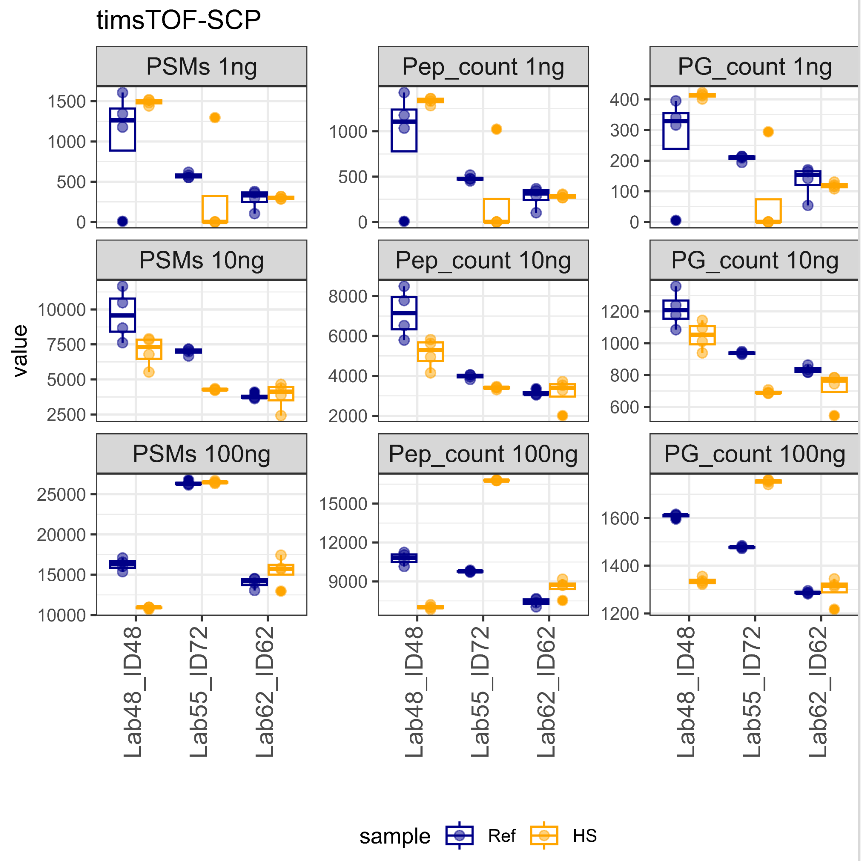


| Dataset | ID48 | ID72 | ID62 |
|---|---|---|---|
| Lab | Lab48 | Lab55 | Lab62 |
| Tube.transfer | 0 | 0 | 0 |
| Reduction | 0 | 0 | 0 |
| Reduction.reagent | NA | NA | NA |
| Alkylation | 0 | 0 | 0 |
| Alkylation.Reagent | NA | NA | NA |
| Enzyme | Tryp | Tryp+LysC | Tryp |
| Enzyme.amount..ng. | 10\|4\|2 | 5 | 10 |
| Digestion.Buffer | 0.2% DDM, TEAB 100mM | 50 mM ABC | 0,2% DDM, 100mM TEAB |
| Volume.of.digestion.buffer..uL. | 4 | 15 | 1 |
| HPLC | U3000 | Bruker nanoElute2 | nanoElute 2 |
| Gradient..min. | 30 | 100 | 115\|77\|77 |
| Column length (cm) | 25 | 25 | 25 |
| MS2.detector | high resolution | HIGH RESOLUTION | High resolution |
| MS_newName | timsTOF-SCP | timsTOF-SCP | timsTOF-SCP |

*ix) ZenoTOF 7600*

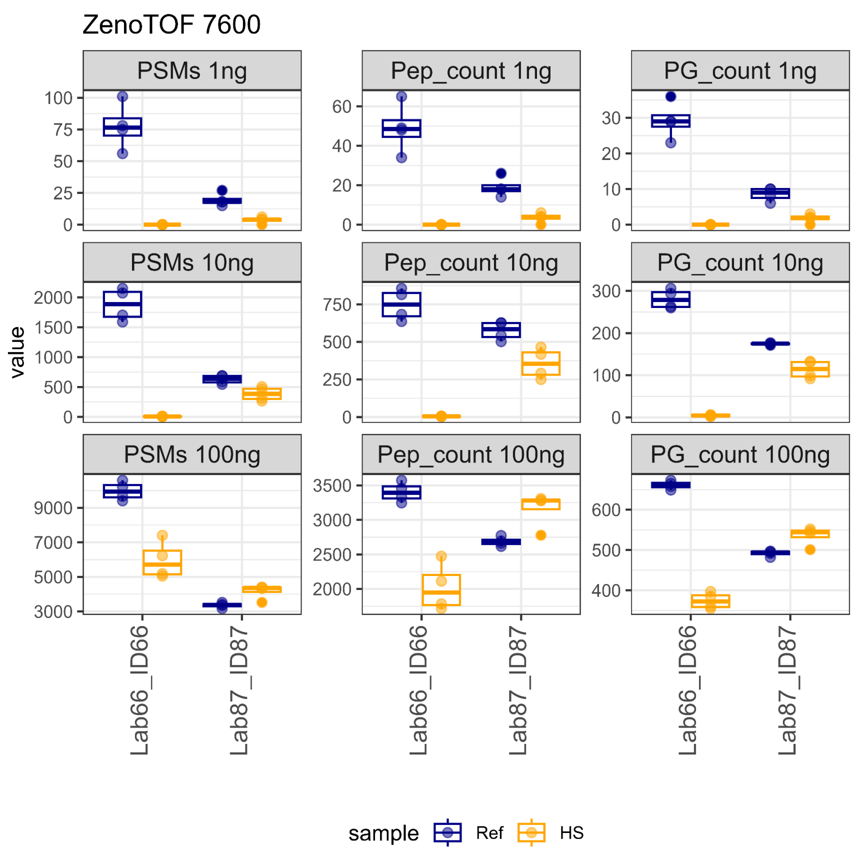


| Dataset | ID66 | ID87 |
|---|---|---|
| Lab | Lab66 | Lab87 |
| Tube.transfer | 1 | 1 |
| Reduction | 0 | 0 |
| Reduction.reagent | NA | NA |
| Alkylation | 0 | 0 |
| Alkylation.Reagent | NA | NA |
| Enzyme | Tryp+LysC | Tryp |
| Enzyme.amount..ng. | 1\|0.1\|0.01 | 25 |
| Digestion.Buffer | TEABC | RapiGest 0.2% |
| Volume.of.digestion.buffer..uL. | 9 | 3 |
| HPLC | Waters nanoAcquity M-class | Evosep One |
| Gradient..min. | 20 | 11 |
| Column length (cm) | 15 | 8 |
| MS2.detector | High resolution TOF | high res |
| MS_newName | ZenoTOF 7600 | ZenoTOF 7600 |

**Figure S4a**: Box plots with the number of identifications (y-axis) for each laboratory and dataset (x-axis) for a given instrument. Grid rows indicate the sample input amounts in ng, and grid columns break down the numbers for Peptide Spectrum Matches (PSMs), peptides, and protein groups (PGs). The color split identification numbers between high sensitivity (HS) and reference (Ref) samples. The table at right shows the collected metadata for each dataset shown. Instruments shown: Exploris 240, Exploris 480, LTQ Orbitrap Velos Pro, Orbitrap Astral, Orbitrap Eclipse, Q Exactive, timsTOF-FLEX, timsTOF-SCP and ZenoTOF 7600.

**b) Outcome metrics analysis breakdown for data independent acquisition (DIA).**

*i) Exploris 480*

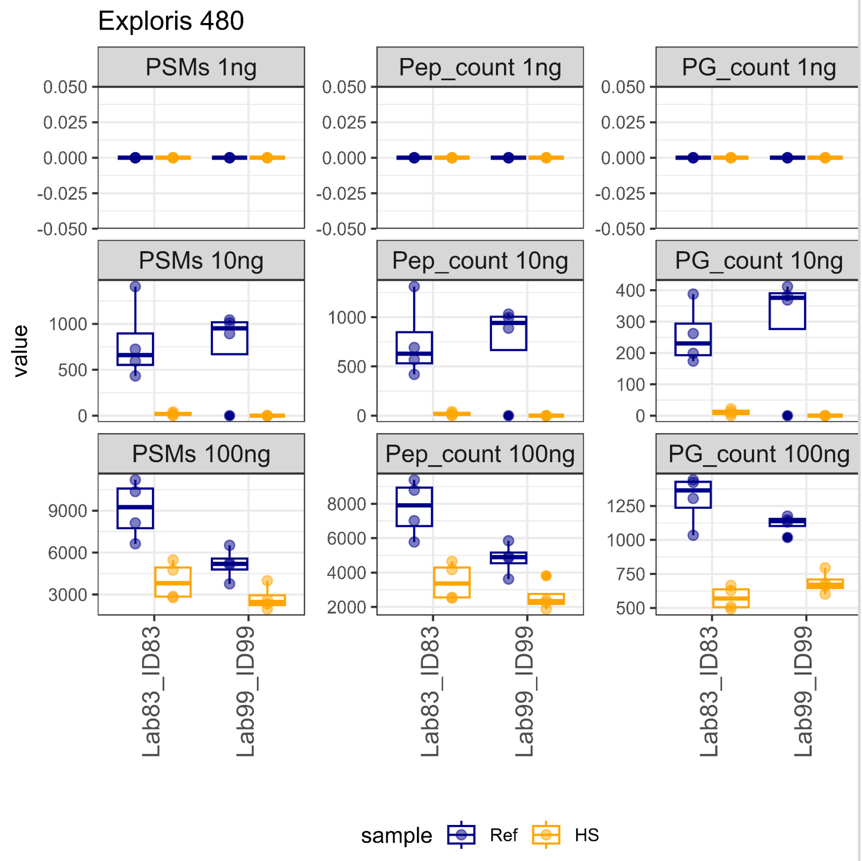


| Dataset | ID83 | ID89 |
|---|---|---|
| Lab | Lab83 | Lab89 |
| Tube.transfer | 0 | 0 |
| Reduction | 1 | 0 |
| Reduction reagent | 4 ÂµL of 100 mM TCEP, 10 mM TCEP final concentration | NA |
| Alkylation | 1 | 0 |
| Alkylation | 4 ÂµL of 400 mM 2-chloroacetamide40 mM 2-chloroacetamide final concentration | NA |
| Reagent | | |
| Enzyme | Tryp+LysC | Tryp |
| Enzyme.amount..ng. | 500 | 20 |
| Digestion.Buffer | 50 mM ABC pH 8.0 | ProteaseMax 0.05%, TEAB 100 mM |
| Volume.of.digestion.buffer..uL. | 20 | 10 |
| HPLC | Vanquish Neo UHPLC | EASY nLC-1200 |
| Gradient..min. | 76 | 65\|20\|20 |
| Column length (cm) | 50 | 25 |
| MS2.detector | | |
| MS_newName | Exploris 480 | Exploris 480 |
| FAIMS | no | yes |

*ii) Orbitrap Astral*

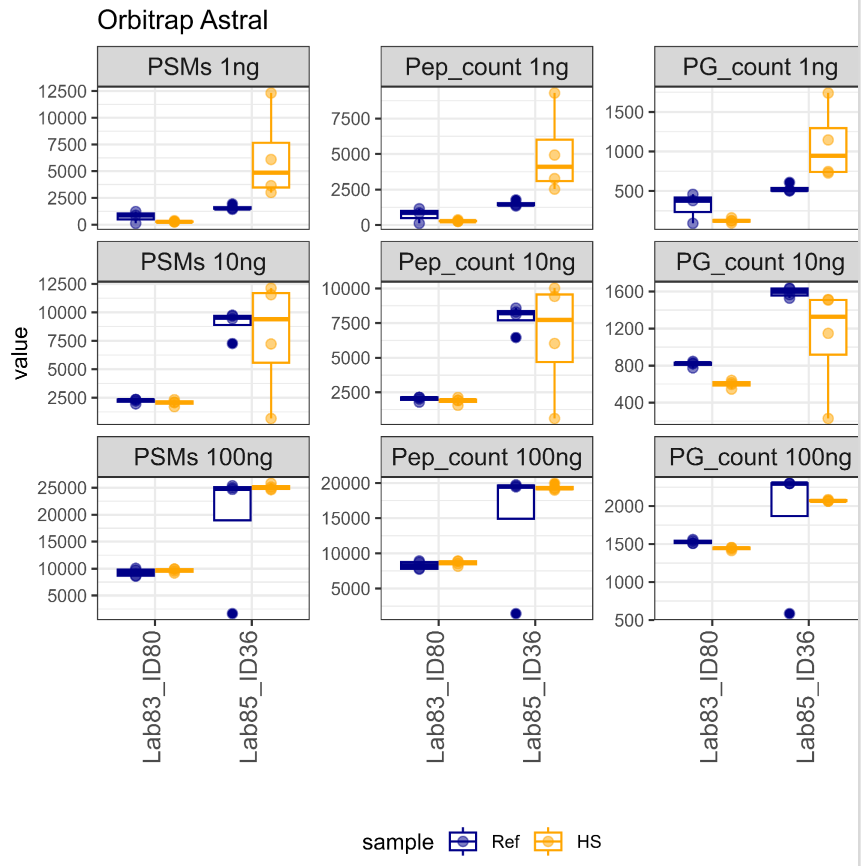


| Dataset | ID36 | ID80 |
|---|---|---|
| **Lab** | **Lab36** | **Lab80** |
| **Tube.transfer** | 1 | 0 |
| **Reduction** | 0 | 0 |
| **Reduction.reagent** | | |
| **Alkylation** | 0 | 0 |
| **Alkylation.Reagent** | | |
| **Enzyme** | Tryp | Tryp+LysC |
| **Enzyme.amount..ng.** | 50 | 50 |
| **Digestion.Buffer** | ABC 0.2M | TEAB 50mM, 0.01% DMM |
| **Volume.of.digestion.buffer..uL.** | 3 | 5 |
| **HPLC** | Vanquish Neo | Vanquish Neo |
| **Gradient..min.** | 30 | 18 |
| **Column length (cm)** | 50 | 5.5 |
| **MS2.detector** | Astral | Astral, 80000 |
| **MS_newName** | Orbitrap Astral | Orbitrap Astral |

*iii) Orbitrap Eclipse*

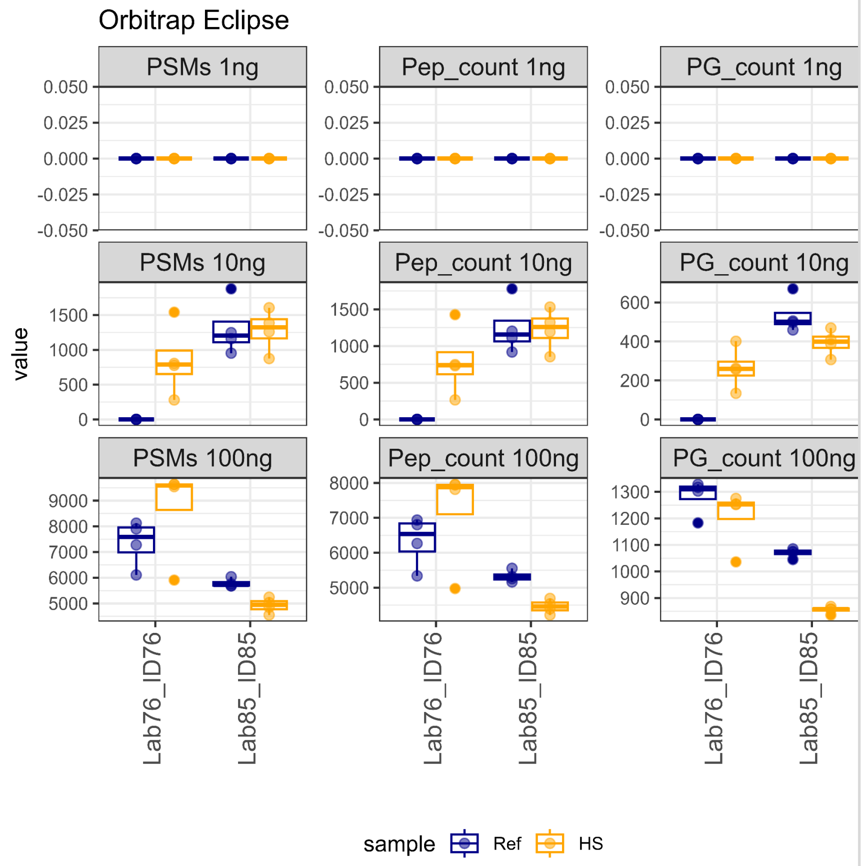


| Dataset | ID76 | ID85 |
|---|---|---|
| Lab | Lab76 | Lab85 |
| Tube.transfer | 1 | 1 |
| Reduction | 0 | 0 |
| Reduction.reagent | NA | NA |
| Alkylation | 0 | 0 |
| Alkylation.Reagent | NA | NA |
| Enzyme | Tryp | Tryp |
| Enzyme.amount..ng. | 75 | 50 |
| Digestion.Buffer | 100mM Ammonium bicarbonate | ABC 0.2M |
| Volume.of.digestion.buffer..uL. | 22 | 3 |
| HPLC | Evosep | Thermo nLC-1200 |
| Gradient..min. | 44 | 60 |
| Column length (cm) | 8 | 50 |
| MS2.detector | Low Res. | ION TRAP |
| Notes | | Inject 90% sample |
| MS_newName | Orbitrap Eclipse | Orbitrap Eclipse |

*iv) timsTOF-FLEX*

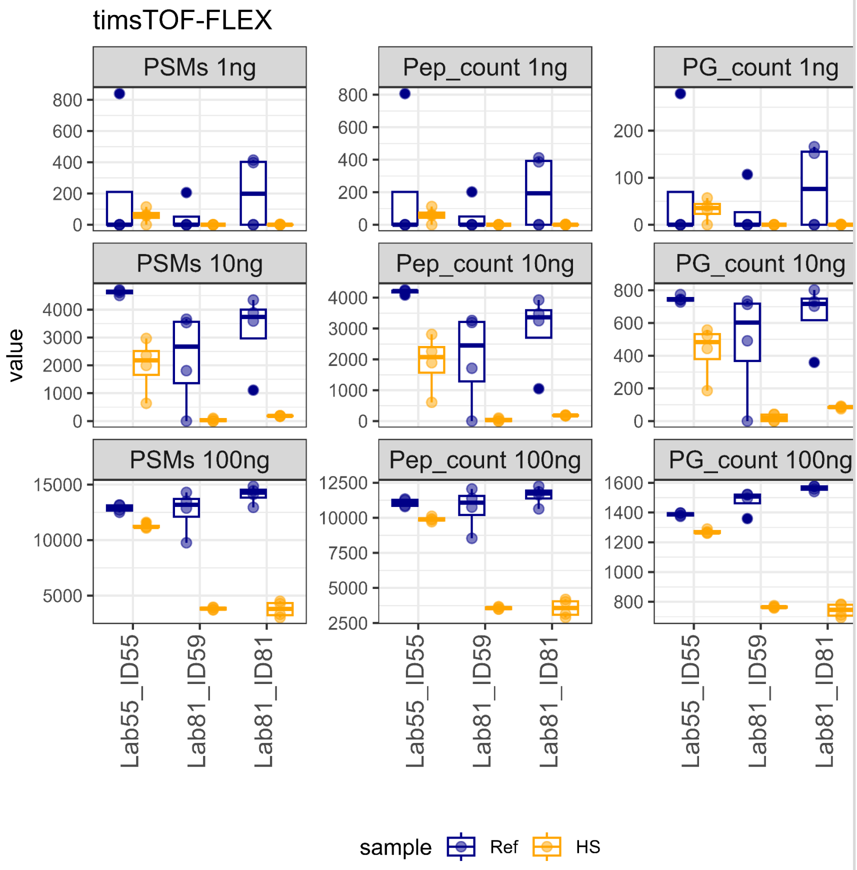


| Dataset | ID55 | ID59 | ID81 |
|---|---|---|---|
| **Lab** | **Lab55** | **Lab81** | **Lab81** |
| **Tube.transfer** | 0 | 1 | 1 |
| **Reduction** | 0 | 1 | 1 |
| **Reduction.reagent** | NA | iSTkit | sp3_iSTkit |
| **Alkylation** | 0 | 1 | 1 |
| **Alkylation.Reagent** | NA | iSTkit | sp3_iSTkit |
| **Enzyme** | Tryp+LysC | Tryp+LysC | Tryp+LysC |
| **Enzyme.amount..ng.** | 5 | NA | NA |
| **Digestion.Buffer** | 50 mM ABC | iSTkit | sp3_iSTkit |
| **Volume.of.digestion.buffer..uL.** | 18 | | |
| **HPLC** | EVOSEP ONE | Evosep | Evosep |
| **Gradient..min.** | 21 | 21 | 21 |
| **Column length (cm)** | 8 | 8 | 8 |
| **MS2.detector** | HIGH RESOLUTION | HIGH | HIGH |
| **Notes** | | iST kits from Preomics to perform the digestions. | sp3 add-on to iST kits from Preomics |
| **MS_newName** | timsTOF-FLEX | timsTOF-FLEX | timsTOF-FLEX |

*v)* *timsTOF-SCP*

**timsTOF-SCP**

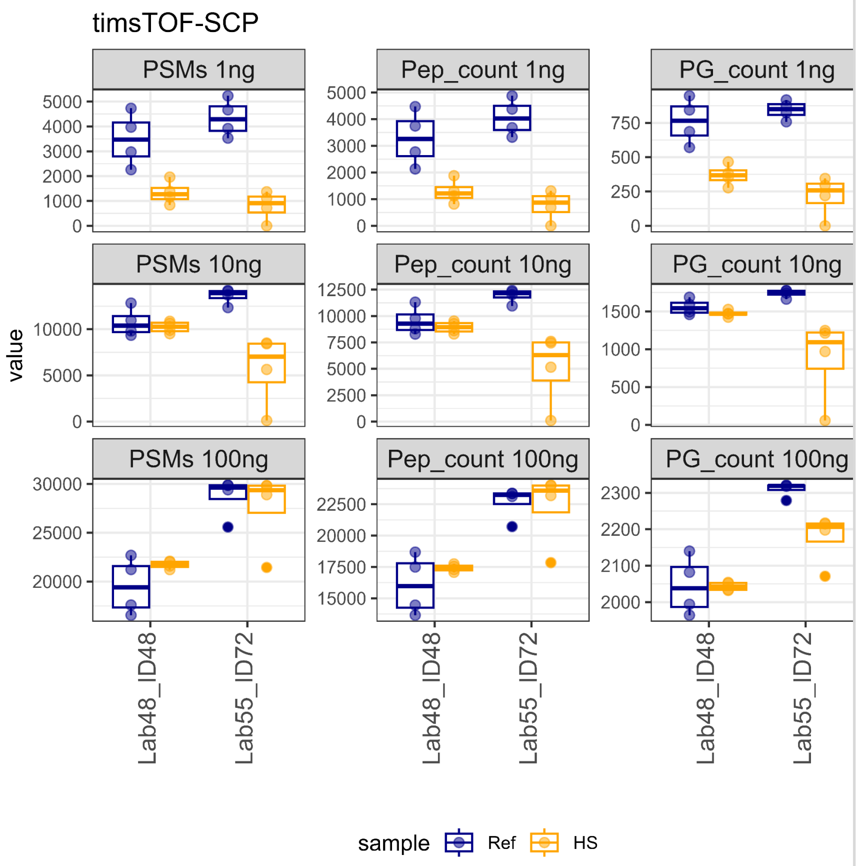


| Dataset | ID48 | ID72 |
|---|---|---|
| **Lab** | **Lab48** | **Lab55** |
| **Tube.transfer** | 0 | 0 |
| **Reduction** | 0 | 0 |
| **Reduction.reagent** | NA | NA |
| **Alkylation** | 0 | 0 |
| **Alkylation.Reagent** | NA | NA |
| **Enzyme** | Tryp | Tryp+LysC |
| **Enzyme.amount..ng.** | 10\|4\|2 | 5 |
| **Digestion.Buffer** | 0.2% DDM, TEAB 100mM | 50 mM ABC |
| **Volume.of.digestion.buffer..uL.** | 4 | 15 |
| **HPLC** | U3000 | Bruker nanoElute2 |
| **Gradient..min.** | 30 | 100 |
| **Column length (cm)** | 25 | 25 |
| **MS2.detector** | high resolution | HIGH RESOLUTION |
| **MS_newName** | timsTOF-SCP | timsTOF-SCP |

*vi) ZenoTOF 7600*

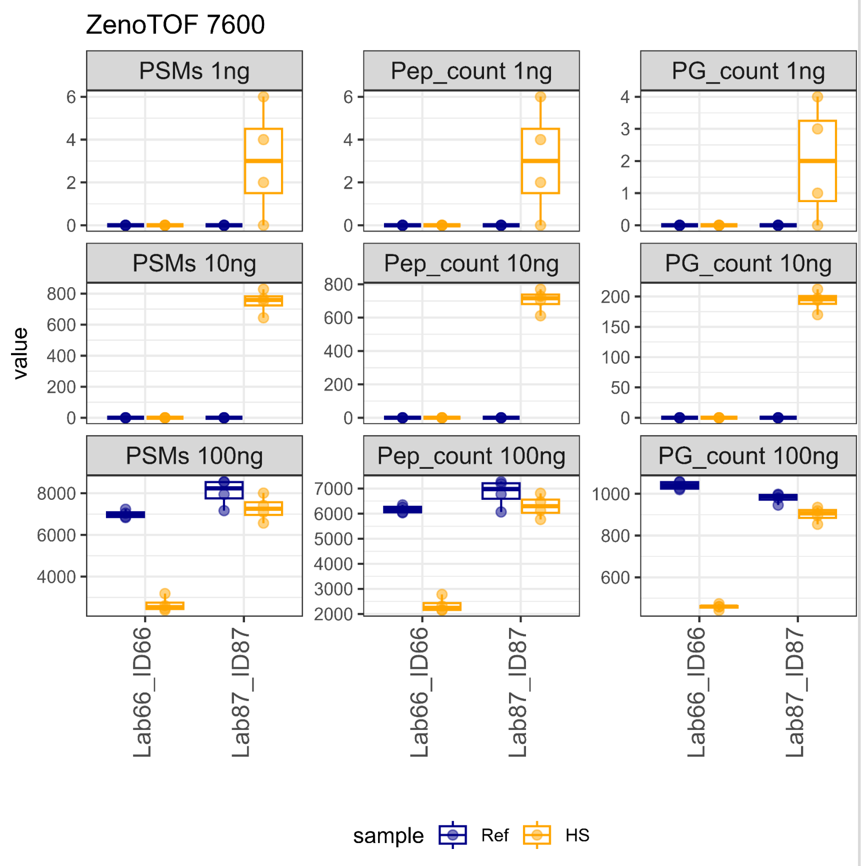


| Dataset | ID66 | ID87 |
|---|---|---|
| Lab | Lab66 | Lab87 |
| Tube.transfer | 1 | 1 |
| Reduction | 0 | 0 |
| Reduction.reagent | NA | NA |
| Alkylation | 0 | 0 |
| Alkylation.Reagent | NA | NA |
| Enzyme | Tryp+LysC | Tryp |
| Enzyme.amount..ng. | 1\|0.1\|0.01 | 25 |
| Digestion.Buffer | TEABC | RapiGest 0.2% |
| Volume.of.digestion.buffer..uL. | 9 | 3 |
| HPLC | Waters nanoAcquity M-class | Evosep One |
| Gradient..min. | 20 | 11 |
| Column length (cm) | 15 | 8 |
| MS2.detector | High resolution TOF | high res |
| MS_newName | ZenoTOF 7600 | ZenoTOF 7600 |

**Figure S4b**: Box plots with the number of identifications (y-axis) for each laboratory and dataset (x-axis) for a given instrument. Grid rows indicate the sample input amounts in ng, and grid columns break down the numbers for Peptide Spectrum Matches (PSMs), peptides, and protein groups (PGs). The color split identification numbers between high sensitivity (HS) and reference (Ref) samples. The table at right shows the collected metadata for each dataset shown. Instruments shown: Exploris 480, Orbitrap Astral, Orbitrap Eclipse, timsTOF-FLEX, timsTOF-SCP and ZenoTOF 7600.
